# Local confinement within plasma membrane nanodomains promotes constitutive activity of GPCRs

**DOI:** 10.64898/2026.02.17.706370

**Authors:** Xiaohan Zhou, Maria Shemeteva, François Simon, Louis-Phillipe Picard, Aiyan Brown, Gurveen Dhillon, Malene L. Urbanus, Lucien E. Weiss, R. Scott Prosser, Claudiu C. Gradinaru

## Abstract

Many G protein-coupled receptors (GPCRs) exhibit constitutive activity through spontaneous conformational transitions that facilitate G protein coupling and downstream signaling in the absence of ligand binding. This baseline activity, which is critical for cellular homeostasis, can be influenced by receptor conformational ensembles and plasma membrane organization. We used live-cell signaling assays, single-particle tracking (SPT) and fluorescence cross-correlation spectroscopy (FCCS) to investigate how membrane organization impacts the basal activity of the M_1_ muscarinic receptor (M_1_R) and the adenosine A_2A_ receptor (A_2A_R). M_1_R showed minimal basal Ca^2+^ responses, whereas A_2A_R exhibited basal cAMP production that was eliminated by an inverse agonist. FCCS showed that in the absence of ligands only a small portion of M_1_R co-diffused with its cognate Gα_11_ protein without ligand, whereas a larger fraction of A_2A_R co-diffused with Gα_s_. Interestingly, SPT revealed that A_2A_R was enriched in slowly diffusing, confined states with spatial scales around 150–200 nm. The sensitivity to cholesterol- and membrane-order-altering agents suggests they are associated with raft-related membrane organization. Dual-color tracking and diffusion mapping demonstrated that A_2A_R and Gα_s_ shared confinement domains under basal conditions, whereas M_1_R and Gα_11_ only showed such co-confinement in the active state. We propose a model in which the co-confinement of GPCRs and G proteins within plasma membrane nanodomains promotes constitutive activity by increasing opportunities for transient productive encounters. This model links receptor conformational dynamics and nanoscale membrane organization to basal signaling output.

**One-sentence summary:** Single-particle tracking experiments show that co-confinement of G protein-coupled receptors and G proteins within membrane nanodomains is an important factor for basal GPCR signaling.

## INTRODUCTION

G protein–coupled receptors (GPCRs) constitute the largest superfamily of membrane signaling proteins and are major targets of clinically approved drugs because of their broad physiological roles and extracellular accessibility (1). In the canonical view of GPCR signaling, agonist binding shifts the receptor conformational ensemble toward active states that engage cognate heterotrimeric G proteins and initiate downstream signaling (**Fig. 1A**) (2). Many GPCRs, however, also signal in the absence of agonist (3). This constitutive, or basal, activity reflects the ability of unliganded receptors to sample signaling-competent conformations and generate downstream responses (**Fig. 1B**). Basal signaling is not simply experimental background, and its magnitude can depend on receptor identity, expression level, G protein availability, and cellular context (4, 5).

**Figure 1.**
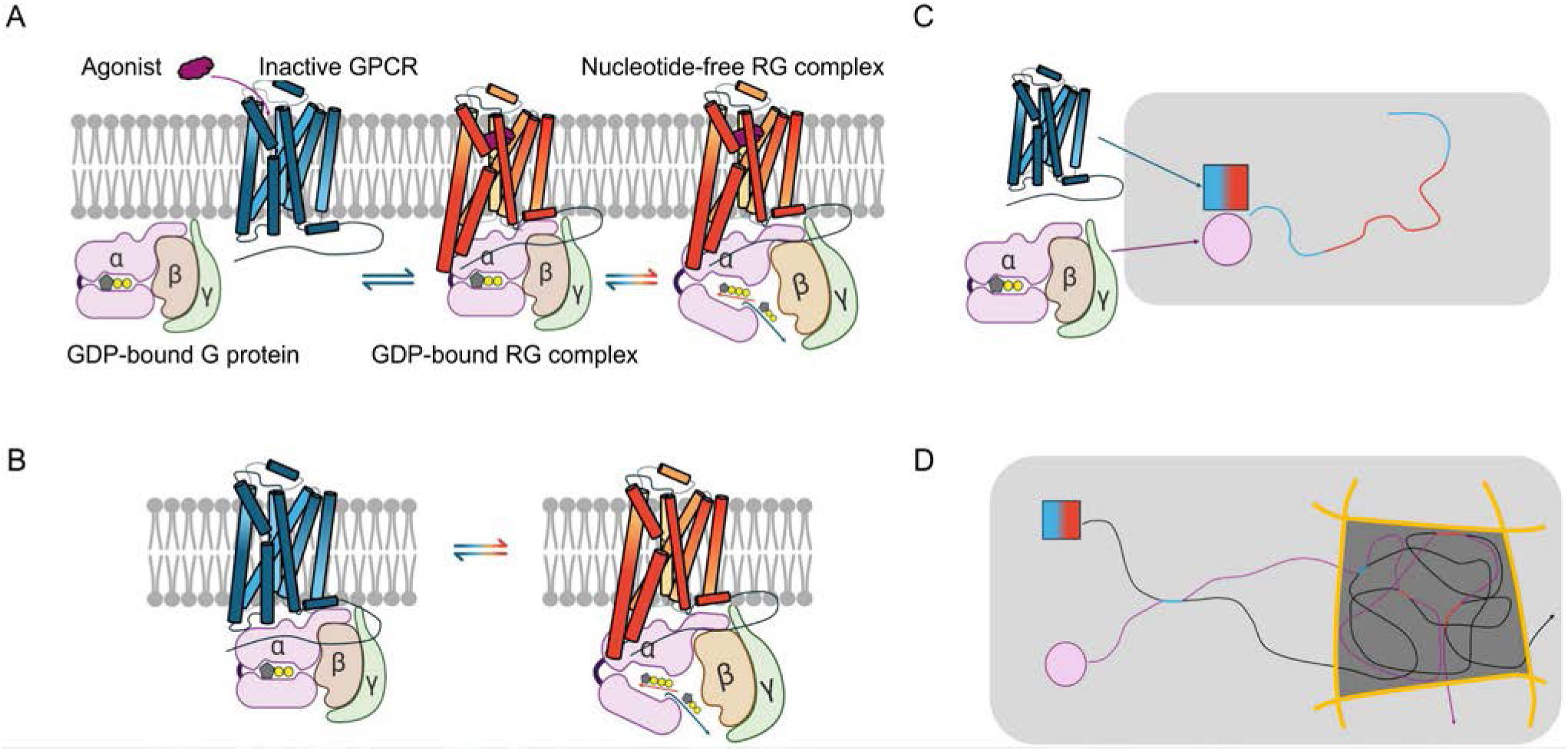
Models of GPCR–G protein coupling and the role of membrane organization in signaling. **(A)** *Canonical activation model*. Agonist binding to inactive GPCR (blue) promotes recruitment of guanosine diphosphate (GDP)-bound G protein to form an RG complex, after which the receptor becomes active (red), drives nucleotide exchange, and triggers *Gβγ* dissociation for downstream signaling. **(B)** *Constitutive signaling model*. In the absence of agonist, receptors spontaneously interconvert between inactive (blue) and active (red) states, couple to G proteins and drive nucleotide exchange. **(C)** *Pre-coupling model*. GPCR and G protein form stable, co-diffusing pre-coupled RG complexes that undergo spontaneous transitions between inactive (blue) and active (red) conformations, producing basal activity. **(D)** *Co-confinement model*. GPCRs (black) and G proteins (violet) diffuse independently and mostly have non-productive encounters (blue, inactive trajectory), but when co-confined within membrane nanodomains, increased local concentration promotes more frequent interactions and transient active (red) RG complexes that enhance basal signaling.

Studies of *β*_2_-adrenergic and *δ*-opioid receptors show substantial basal activity that can be suppressed by inverse agonists (6, 7), and broader surveys reveal large differences in basal signaling across GPCR families (4). At one extreme, rhodopsin exhibits little detectable activity in the dark state (8, 9), whereas the histamine H_3_ receptor can reach 40–60% of its maximal agonist-induced response without ligand (10). Orphan GPCRs, for which natural endogenous ligands remain unclear, often exhibit high basal activity (11). An entire subclass of adhesion GPCRs appear to have built-in activity – for example, GPR64, GPR114 and GPR126 all display constitutive signaling functions due to their tethered agonism (12) A study on GPR35 further shows pathway-selective constitutive Gα_12/13_ signaling that contributes to epithelial barrier regulation (13).

At the molecular level, constitutive activity is commonly described in terms of receptor conformational equilibria. Extended ternary-complex and cubic models incorporate ligand binding and G protein association into this framework, allowing receptor–G protein (RG) complexes to form even in the absence of agonist (5). This framework also accommodates pre-coupling, in which receptors and G proteins remain spatially associated before activation and signaling proceeds through conformational rearrangement within an existing complex (**Fig. 1C**). Proximity-based assays, including fluorescence resonance energy transfer (FRET) and bioluminescence resonance energy transfer (BRET), provide evidence for such preassembled complexes, including *β*_2_AR–Gα_s_ and *α*_2*A*_-adrenergic receptor (α_2A_AR)–Gα_o_ complexes (14, 15). Structural studies further support the idea that receptor–G protein association is not restricted to agonist-bound active states. Cryogenic electron microscopy revealed *κ*-opioid receptor (KOR)–G_i_ complexes bound to inverse agonists, showing that inverse agonism can occur within RG complexes rather than simply by preventing G protein binding (16).

Stable pre-coupling, however, is unlikely to account for all receptor–G protein communication. An alternative model is that receptors and G proteins diffuse independently and interact through repeated transient encounters. If both partners spend sufficient time within the same membrane compartment, their local concentration and probability of re-encounter can increase without requiring a long-lived preassembled complex (**Fig. 1D**). In this model, productive RG coupling depends not only on receptor conformation and G protein availability, but also on how efficiently the two partners gain spatial access to one another.

The spatial component is important because the plasma membrane cannot be treated simply as a homogeneous two-dimensional fluid. Receptors, G proteins, and other membrane components can become transiently restricted within membrane compartments, altering their local mobility, residence time, and encounter probability. Confined motion can arise from several physical mechanisms, including transient binding, receptor clustering, cytoskeletal barriers, protein scaffolds, and lipid-mediated heterogeneity (17, 18). In the picket-and-fence model, cortical actin forms submembrane fences and actin-anchored transmembrane proteins act as pickets that partition the membrane into dynamic compartments and restrict long-range diffusion (17, 19). In parallel, preferential lipid–lipid and lipid–protein interactions can generate compositional heterogeneity within the bilayer. Cholesterol and sphingolipids can promote small, heterogeneous, dynamic liquid-ordered (Lo)-like nanodomains that transiently enrich or exclude particular membrane components, forming the basis of the modern lipid raft concept (20, 21). These mechanisms can coexist and influence one another rather than representing separate models of membrane organization. Importantly, confined diffusion describes molecular motion but does not by itself identify the physical origin of the compartment. A confinement size comparable to reported Lo-like nanodomains, or sensitivity to cholesterol- and membrane-modifying treatments, is consistent with lipid-mediated organization but does not directly establish that a confined region is a lipid raft.

The same distinction applies to GPCR signaling hotspots. Single-molecule imaging shows that receptor–G protein interactions can occur preferentially within restricted regions of the plasma membrane rather than uniformly across the cell surface (22). Such hotspots can be viewed as dynamic regions in which local confinement or molecular enrichment increases the probability of receptor–effector encounters, but the term does not specify a unique membrane structure or composition. An additional molecular reference is needed to test whether these compartments are associated with raft-related membrane organization. CD59 is a glycosylphosphatidylinositol (GPI)-anchored outer-leaflet protein widely used in studies of raft-associated nanoscale organization. Single-molecule measurements show that GPI-anchored proteins, including CD59, can participate in transient cholesterol-dependent assemblies rather than stable membrane islands (23). CD59 thus provides a useful raft-associated marker for asking whether GPCR-containing membrane compartments preferentially overlap with regions exhibiting similar organization, without assigning their lipid identity from diffusion behavior alone.

Here, we investigated these questions using the M_1_ muscarinic receptor (M_1_R) and the adenosine A_2A_ receptor (A_2A_R), two class A GPCRs that play pivotal roles in the central nervous system and have markedly different basal signaling properties. M_1_R, which is expressed in cortical and hippocampal neurons, couples predominantly to Gα_q/11_ and exhibits low constitutive activity in heterologous systems unless Gα_q_ is strongly overexpressed (24). In contrast, A_2A_R, which is abundant in the basal ganglia, couples to Gα_s/o_ and exhibits substantial agonist-independent cyclic adenosine monophosphate (cAMP) production (25). Previous studies further show that A_2A_R exchanges between distinct mobility states and that access to slowly diffusing membrane compartments depends on cholesterol, the receptor C terminus, and associated cellular components (26, 27). These receptors provide complementary systems for asking whether differences in basal signaling are accompanied by differences in receptor–G protein organization within the plasma membrane.

We combined live-cell signaling assays with single-molecule fluorescence imaging and spectroscopy to relate signaling output to receptor and G protein mobility and spatial organization. We first asked whether basal activity is accompanied by receptor–G protein co-diffusion and shared confinement within membrane compartments, which could increase opportunities for repeated local encounters. We then characterized CD59 diffusion and quantified its co-confinement with M_1_R and A_2A_R under basal and agonist-stimulated conditions to test whether these receptor-containing compartments are associated with raft-related membrane organization. The results lead to a model in which constitutive GPCR activity emerges from two spatiotemporally linked factors: the ability of a receptor to sample signaling-competent conformations and its spatial access to the appropriate G protein within the heterogeneous plasma membrane.

## RESULTS

### Signaling assays reveal different levels of constitutive activity for M_1_R vs A_2A_R

The constitutive activity of GPCRs was studied in Flp-In T-REx 293 cell lines stably expressing Halo-A_2A_R or Halo-M_1_R (28). These cell lines support inducible, controlled receptor expression (29). The basal activity and functionality of the receptors were quantified through their downstream secondary messenger responses in the absence and presence of orthosteric ligands. In Halo-M_1_R-expressing cells, the full agonist carbachol increased the Ca^2+^ response in a concentration-dependent manner compared with vehicle treatment (**Fig. 2A**). The estimated half-maximal effective concentration (*EC*_50_) of 5.8 ± 1.9 *μM* was similar to reported values for wild-type M_1_R (30), supporting the functionality of the expressed Halo-M_1_R. However, the inverse agonist pirenzepine did not significantly change the Ca^2+^ response compared with vehicle treatment, indicating very low (undetectable) basal muscarinic Gα_q/11_–Ca^2+^ signaling under our conditions. Because human embryonic kidney 293 (HEK293)-derived cells can express endogenous M_3_ muscarinic receptors (M_3_R), some contribution to the bulk Ca^2+^ response cannot be excluded.

**Figure 2.**
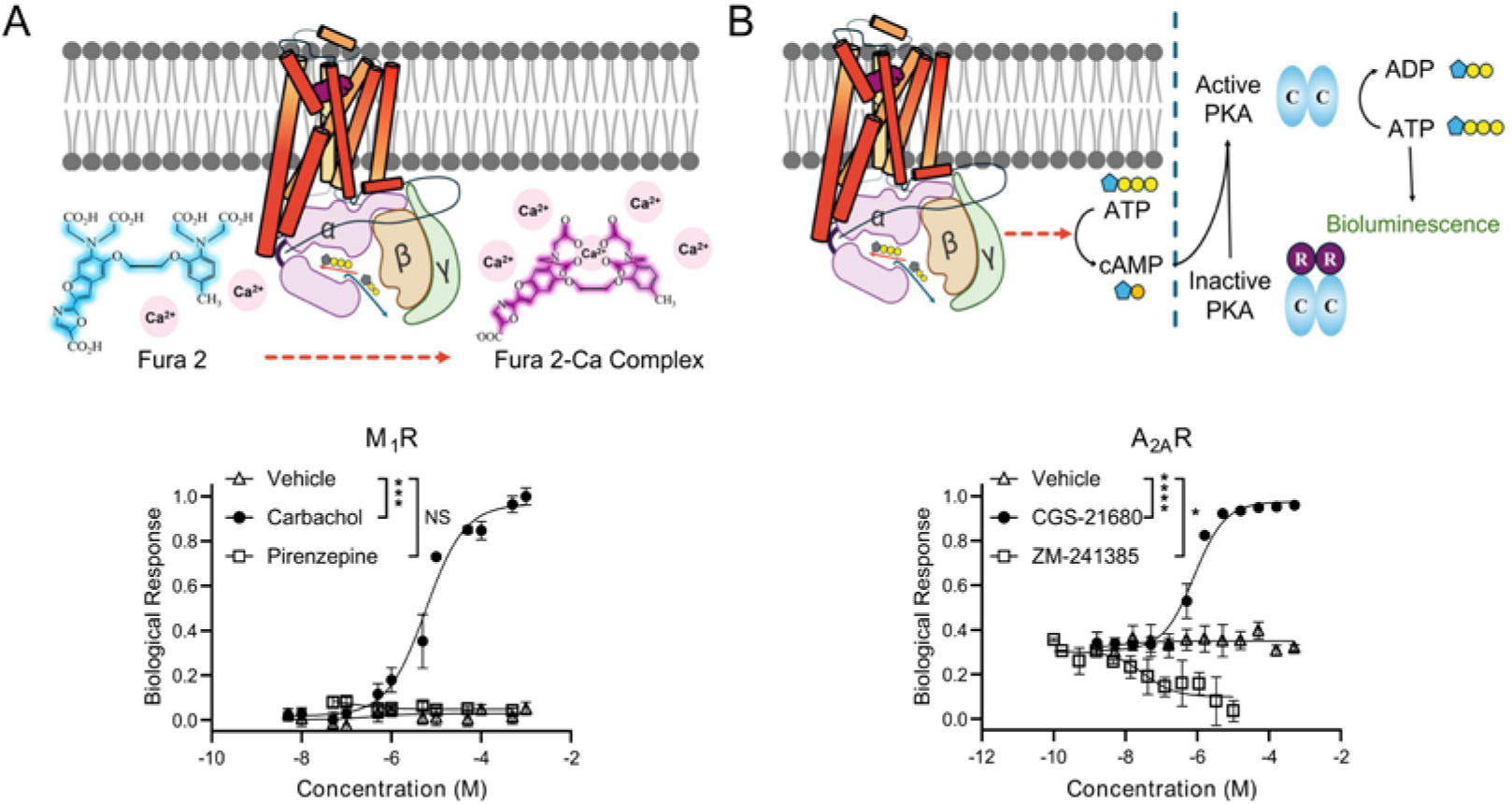
Cell signaling assays. **(A)** In Flp-In T-REx 293 cells stably expressing Halo-M_1_R, the intracellular Ca^2+^ concentration was measured by spectral changes of the Fura-2 dye after treatment of live cells with agonist (*carbachol*) or inverse agonist (*pirenzepine*), using no ligand (DMSO vehicle) as control. **(B)** In Flp-In T-REx 293 cells stably expressing Halo-A_2A_R, changes in cAMP concentration were measured by luciferin bioluminescence mediated through adenosine triphosphate (ATP) hydrolysis by protein kinase A (PKA), after 30 minutes treatment of cells with agonist (*CGS-21680*) or inverse agonist (*ZM-241385*), with DMSO vehicle as control. Data in **(A)** and **(B)** are means ± SD from *n* = 3 independent biological experiments per condition/concentration. Raw responses were averaged at each concentration and fitted by nonlinear regression using a four-parameter logistic model; *EC*_50_ and half-maximal inhibitory concentration (*IC*_50_) values are reported with 95% confidence intervals. Data were subsequently normalized between 0 and 1 for presentation. *E*_max_ values were compared between each treatment and its vehicle control using two-sided Welch’s *t*-tests: *, *P* < 0.05; ***, *P* < 0.001; ****, *P* < 0.0001; NS, not significant (*P* ≥ 0.05).

In Halo-A_2A_R-expressing cells, the full agonist CGS-21680 increased cAMP production through the Gα_s_ pathway, whereas the inverse agonist ZM-241385 reduced it in a concentration-dependent manner compared with vehicle treatment (**Fig. 2B**). These observations indicate measurable tonic activity in the apo state (about 25% of maximum response), consistent with previous radiolabeled cAMP formation assays (25). The agonist potencies for both GPCRs were comparable to reported values (30), confirming the functionality of our protein constructs.

### FCCS: different levels of co-diffusion for M_1_R vs A_2A_R with their cognate G proteins in the basal state

To relate constitutive GPCR activity to receptor–G protein co-diffusion, we used fluorescence cross-correlation spectroscopy (FCCS) (31) in cells co-expressing Halo-M_1_R and Gα_11_-SNAP, or Halo-A_2A_R and Gα_s_-SNAP. Calibration measurements and the live-cell FCCS positive control are shown in **fig. S1A–D**. In the absence of ligands, M_1_R showed a comparatively uniform plasma-membrane distribution, whereas A_2A_R displayed more pronounced spatial heterogeneity with local receptor-enriched regions (**Fig. 3A,B**). The bright cell edges reflect higher apparent fluorescence at the lateral plasma membrane, where more membrane was sampled within the confocal volume. The bright central signal mainly reflects intracellular/internalized receptor rather than plasma-membrane clustering (**Fig. 3A**).

**Figure 3.**
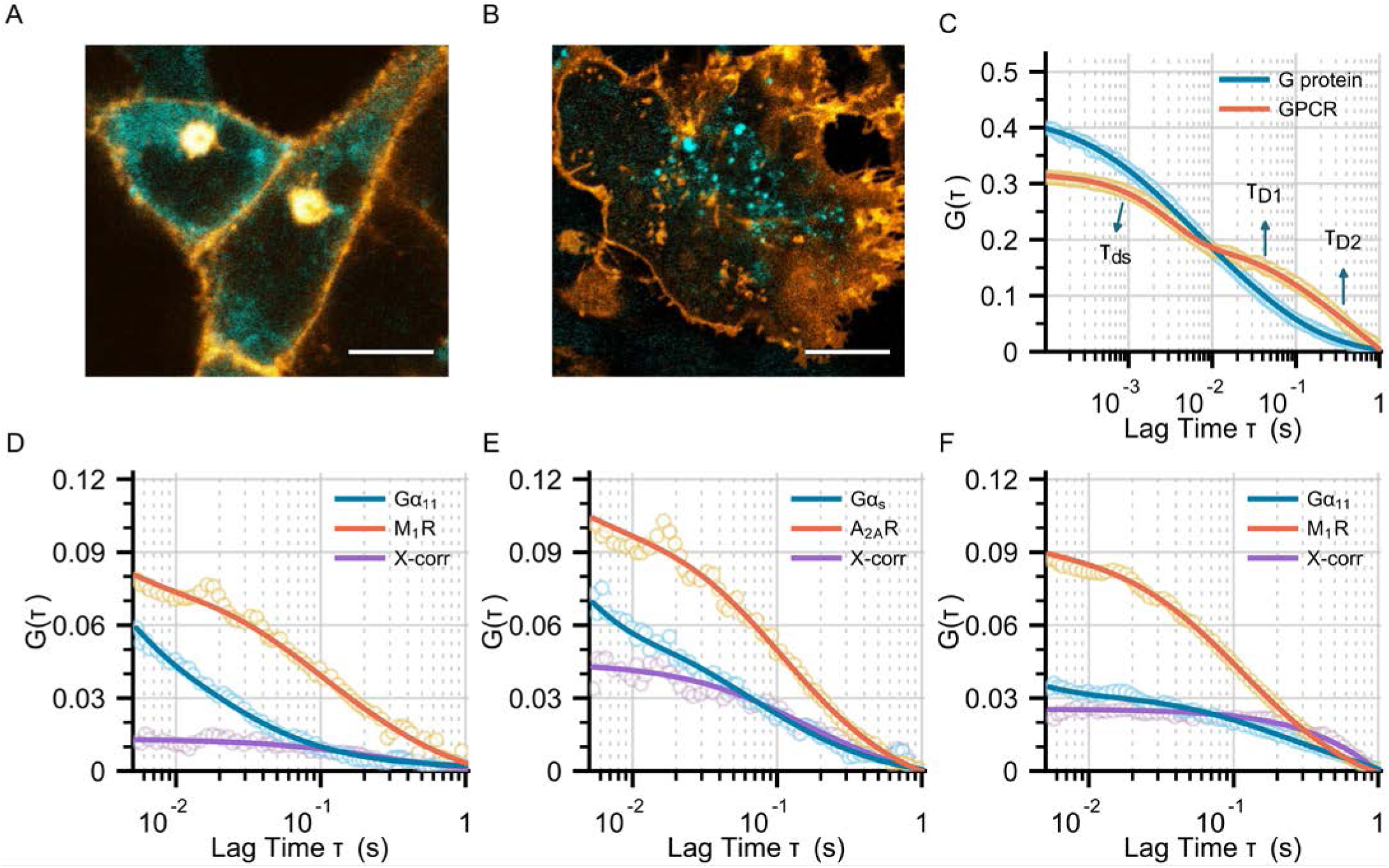
Co-localization and co-diffusion of GPCRs and G proteins in live cells. Representative confocal images at the bottom membrane of live cells co-expressing M_1_R (orange) and Gα_11_ (cyan) **(A)**, or A_2A_R (orange) and Gα_s_ (cyan) **(B)**; scale bar: 5 *μm*. **(C)** Representative autocorrelation (AC) curves for A_2A_R (orange) and Gα_s_ (cyan), highlighting decays corresponding to diffusion (*τ_D_*) and dark-state photophysics (*τ_ds_*). Basal-state (no ligand) dual-color cross-correlation (X-corr) curves for M_1_R and Gα_11_ **(D)** or A_2A_R and Gα_s_ **(E)**; orange, AC of receptors; cyan, AC of G proteins; violet, X-corr between receptors and G proteins. **(F)** Active-state dual-color correlation data for M_1_R and Gα_11_ in the presence of 10*μ*M carbachol. See **Table 1** and **Table 2** for the full list of one- and two-color correlation fitting parameters. Single-color FCS used stable Flp-In T-REx 293 lines, with *n* = 8 A_2A_R and 7 Gα_s_ cells in the datasets illustrated in **(C)**. Dual-color FCCS used stable Gα_11_-SNAP or Gα_s_-SNAP Flp-In T-REx 293 cells transiently expressing the cognate Halo-GPCR. The datasets represented in **(A,D)**, **(B,E)**, and **(F)** comprised *n* = 16, 14, and 13 cells, respectively. Each dataset was collected across three independent biological replicates.

**Table 1.** Single-color FCS fitting parameters^a^.

| Protein | Density (molec./$\mu\text{m}^2$) | $D_1$ ($\mu\text{m}^2/\text{s}$) | $f_1$ | $D_2$ ($\mu\text{m}^2/\text{s}$) | $n$ (cells) |
| --- | --- | --- | --- | --- | --- |
| M<sub>1</sub>R | $4.6 \pm 0.4$ | $0.24 \pm 0.05$ | $76 \pm 9\%$ | $0.02 \pm 0.002$ | 10 |
| A<sub>2A</sub>R | $10.5 \pm 0.8$ | $0.23 \pm 0.002$ | $59 \pm 7\%$ | $0.02 \pm 0.002$ | 8 |
| G$\alpha_{11}$ | $17.6 \pm 0.2$ | $0.63 \pm 0.04$ | 100% | – | 7 |
| G$\alpha_s$ | $16.5 \pm 0.3$ | $0.49 \pm 0.04$ | 100% | – | 7 |

<sup>a</sup> Single-color autocorrelation (AC) data fitted to **Eq. S4** for GPCRs or **Eq. S5** for G proteins. For each experimental condition, the mean values obtained from the global fit are listed, with errors reported as ± SEM.

**Table 2.** Dual-color FCCS fitting results^a^.

| Protein pair | Condition | Density (molec./$\mu\text{m}^2$) | $D_1$ ($\mu\text{m}^2/\text{s}$) | $f_1$ (%) | $D_2$ ($\mu\text{m}^2/\text{s}$) | $fcd$ | $n$ (cells) |
| --- | --- | --- | --- | --- | --- | --- | --- |
| M<sub>1</sub>R + G$\alpha_{11}$ | Apo (R) | 18.4 ± 3.2 | 0.25 ± 0.01 | 87 ± 4 | 0.08 ± 0.01 | 0.27 ± 0.03 | 16 |
| | Apo (G) | 91.8 ± 16.1 | 0.51 ± 0.12 | 94 ± 3 | 0.08 ± 0.01 | 0.11 ± 0.02 | 16 |
| M<sub>1</sub>R + G$\alpha_{11}$ | Active (R) | 28.5 ± 6.5 | 0.21 ± 0.02 | 86 ± 3 | 0.02 ± 0.005 | 0.67 ± 0.07 | 13 |
| | Active (G) | 104 ± 24 | 0.36 ± 0.09 | 33 ± 1 | 0.02 ± 0.005 | 0.43 ± 0.07 | 13 |
| A<sub>2A</sub>R + G$\alpha_s$ | Apo (R) | 22.2 ± 3.1 | 0.23 ± 0.01 | 82 ± 5 | 0.11 ± 0.01 | 0.74 ± 0.05 | 14 |
| | Apo (G) | 74.3 ± 15.6 | 0.41 ± 0.05 | 76 ± 3 | 0.11 ± 0.01 | 0.52 ± 0.06 | 14 |

<sup>a</sup> Dual-color FCCS autocorrelation (AC) data was fitted globally to **Eq. S4**, and cross-correlation (X-corr) data was fitted to **Eq. S6**; $fcd$ was calculated using **Eq. S7**. For each experimental condition, the mean values obtained from the global fit are listed, with errors reported as ± SEM.

Single-color fluorescence correlation spectroscopy (FCS) autocorrelation (AC) measurements in cell lines expressing each receptor or G protein separately revealed their individual membrane diffusion behavior (**Fig. 3C**). The receptor AC curves contained two diffusion components and a photophysical dark-state component. The dark-state lifetime of 2.5 *ms* was much longer than the microsecond-scale triplet-state lifetime of rhodamine-based Janelia Fluor (JF) dyes (32). Single-molecule blinking in fixed Halo-A_2A_R-expressing cells occurred on the same low-millisecond timescale as the slow dark-state kinetics captured in the AC data (**fig. S2A–C**).

Global analysis showed that a large fraction of M_1_R (∼ 75%) diffused fast (*D*_1_ ≈ 0.25 *μm*^2^/*s*), whereas the rest (∼ 25%) exhibited much slower diffusion (*D*_2_ ≈ 0.02 *μm*^2^/*s*). For A_2A_R, the slow-diffusing fraction was higher (∼ 40%) at the expense of the fast-diffusing fraction (∼ 60%) (**Table 1**), consistent with its more localized receptor-rich regions (**Fig. 3B**). In contrast, a single diffusion component was sufficient to describe the G protein AC curves (**Fig. 3C**). Both Gα_11_ and Gα_s_ diffused faster than the receptors (*D* = 0.5–0.6 *μm*^2^/*s*), as expected for membrane-anchored *vs* transmembrane proteins, and their diffusion was comparable to reported values for G proteins and other membrane-anchored proteins (22, 33, 34).

FCCS measurements in cells co-expressing the cognate receptor–G protein pairs revealed co-diffusion of each receptor with its binding partner (**Fig. 3D–F**). The receptor and G protein AC curves shared a slow diffusion component with the cross-correlation (CC) curve, providing an estimate of the co-diffusing fraction (*fcd*).

Without ligand, M_1_R and Gα_11_ exhibited low CC amplitude, indicating that only a minor fraction of the receptor (*fcd_r_* ≈ 30%) co-diffused with Gα_11_ (**Fig. 3D**; **Table 2**). Gα_11_ diffusion was only slightly slower in the presence of M_1_R (**Table 2**). Carbachol stimulation increased the CC amplitude, with up to ∼ 70% of the receptor co-diffusing with Gα_11_ (**Fig. 3F**; **Table 2**). Under this condition, over 60% of Gα_11_ underwent slow diffusion at a rate similar to the slow component in receptor-only cells (*D*_2_ ≈ 0.02 *μm*^2^/*s*).

In contrast, A_2A_R and Gα_s_ showed high CC amplitude even without ligand, indicating that ∼ 75% of the receptors co-diffused with G proteins in the cell membrane (**Fig. 3E**; **Table 2**). The slow-diffusing Gα_s_ population was approximately 25%, with *D*_2_ ≈ 0.11 *μm*^2^/*s*, intermediate between the fast and slow receptor-only components, consistent with a mixture of those populations (**Table 2**). Across FCCS measurements, the fraction of G proteins co-diffusing with receptors (*fcd_gg_*) was lower than the fraction of receptors co-diffusing with G proteins (*fcd_r_*), consistent with the higher local membrane density of G proteins (**Table 2**). Additional apo-state measurements across regions of different receptor density showed density-dependent variation in the co-diffusing fraction, particularly for A_2A_R–Gα_s_ (**fig. S3A–C**).

### Mapping the diffusion of receptors and G proteins at single-molecule level

To examine the physical origin of receptor–G protein co-diffusion, we used single-particle tracking (SPT) in cells expressing Halo-M_1_R or Halo-A_2A_R (**Fig. 4A,B**). Receptor diffusion was compared under basal conditions, after agonist stimulation, and after membrane perturbation. *ExaTrack* analysis (35) distinguished Brownian, confined, and immobile motion. Regularized Bayesian information criterion (BIC) analysis and inspection of the fitted states supported this three-state representation (**fig. S4C**). Although the four-state G protein fit had a lower BIC_reg_, the additional state mainly subdivided the Brownian population.

**Figure 4.**
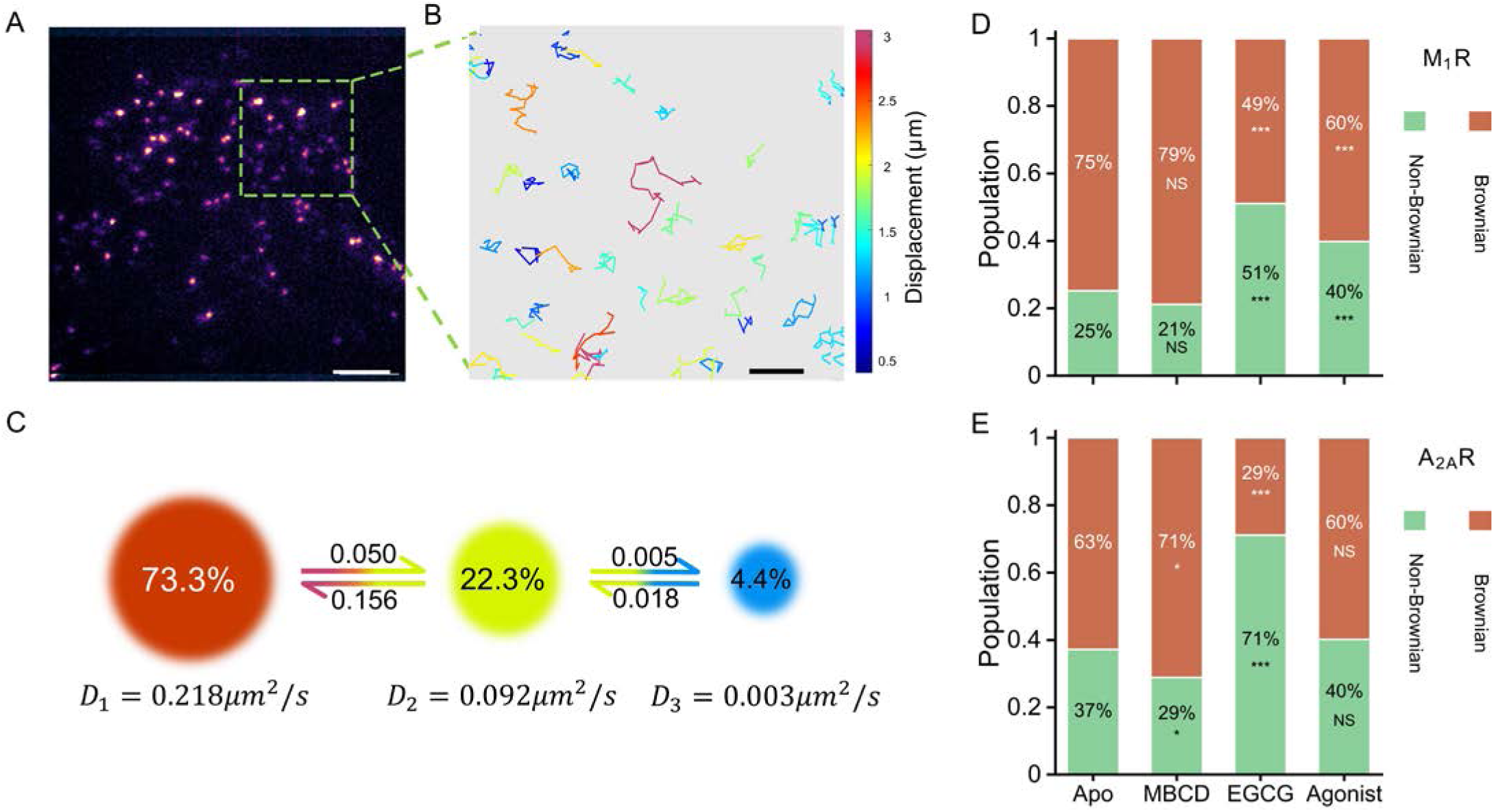
Tracking the diffusion of M_1_R and A_2A_R in the membrane of live cells. SPT used Flp-In T-REx 293 cells: *n* = 1 cell co-expressing A_2A_R and Gα_s_ for **(A–C)**, and stable Halo-M_1_R or Halo-A_2A_R lines for **(D,E)**. Representative particle locations were detected in each frame **(A)** then linked to build 2D diffusion trajectories **(B)**, and color-coded according to their displacement; scale bar: 5 *μm* for **(A)** and 1 *μm* for **(B)**. **(C)** In the example shown, global analysis of all trajectories using ExaTrack identified three interchanging diffusion states: *free* (*D*_1_ ∼ 0.2 *μm*^2^/*s*), *confined* (*D*_2_ = 0.05–0.10 *μm*^2^/*s*), and *immobile* (*D*_3_ < 0.005 *μm*^2^/*s*). Stacked composition bars summarize the mean diffusion populations of M_1_R **(D)** and A_2A_R **(E)** as non-Brownian states (confined plus immobile) and Brownian diffusion, and show changes depending on lipid environment and activation state relative to apo state (*, *P* < 0.05; **, *P* < 0.01; ***, *P* < 0.005; NS, not significantly different, tested by two-sided Welch’s *t*-tests with Holm correction). Full results of SPT analysis, including means ± standard error of the mean (SEM), are listed in **Table 3** and **table S1**. **(D)** *n* = 25, 12, 14, and 15 cells for Apo, MBCD, EGCG, and carbachol, respectively. **(E)** *n* = 23, 11, 15, and 12 cells for Apo, MBCD, EGCG, and CGS-21680, respectively. Data were collected across three independent biological replicates per condition.

**Table 3.** SPT analysis for GPCRs and G proteins under different conditions^a^.

| Protein | Condition | $D_{Br}$ ($\mu m^2/s$) | $D_{conf}$ ($\mu m^2/s$) | $f_{NBr}$ (%)<sup>b</sup> | $R_{conf}$ (nm)<sup>c</sup> | $n$ (cells) |
| --- | --- | --- | --- | --- | --- | --- |
| M<sub>1</sub>R | Apo | $0.218 \pm 0.024$ | $0.092 \pm 0.006$ | $25.2 \pm 5.4$ | $160 \pm 5$ | 25 |
| | + MBCD | $0.174 \pm 0.012$ | $0.054 \pm 0.008$ | $21.1 \pm 2.3$ | $226 \pm 17$ | 12 |
| | + EGCG | $0.178 \pm 0.010$ | $0.028 \pm 0.002$ | $51.1 \pm 7.3$ | $124 \pm 5$ | 14 |
| | + Carbachol | $0.215 \pm 0.016$ | $0.064 \pm 0.006$ | $39.8 \pm 2.4$ | $226 \pm 11$ | 15 |
| G$\alpha_{11}$ | Apo | $0.547 \pm 0.034$ | $0.059 \pm 0.007$ | $36.2 \pm 8.7$ | $141 \pm 8$ | 15 |
| | + Carbachol | $0.626 \pm 0.049$ | $0.120 \pm 0.015$ | $52.1 \pm 6.1$ | $256 \pm 16$ | 13 |
| A<sub>2A</sub>R | Apo | $0.208 \pm 0.012$ | $0.125 \pm 0.009$ | $37.2 \pm 4.8$ | $168 \pm 6$ | 23 |
| | + MBCD | $0.170 \pm 0.013$ | $0.053 \pm 0.003$ | $28.9 \pm 1.7$ | $166 \pm 7$ | 11 |
| | + EGCG | $0.149 \pm 0.008$ | $0.020 \pm 0.002$ | $71.1 \pm 9.1$ | $94 \pm 5$ | 15 |
| | + CGS-21680 | $0.214 \pm 0.015$ | $0.077 \pm 0.008$ | $40.2 \pm 3.4$ | $133 \pm 7$ | 12 |
| G$\alpha_s$ | Apo | $0.449 \pm 0.023$ | $0.088 \pm 0.010$ | $49.6 \pm 7.3$ | $162 \pm 9$ | 18 |
| | + CGS-21680 | $0.467 \pm 0.037$ | $0.064 \pm 0.006$ | $46.3 \pm 6.3$ | $193 \pm 9$ | 13 |

<sup>a</sup> SPT data was analyzed using ExaTrack (35); for each condition, the weighted mean values across cells are listed, with errors reported as ± SEM. <sup>b</sup> $f_{NBr}$ , percentage of the non-Brownian population, calculated as the sum of the confined and immobile fractions. <sup>c</sup> $R_{conf}$ , confinement radius for the confined diffusion population, calculated using **Eq. S14**.

The three-state model characterized diffusion coefficients, state occupancies, and transitions between states (**Fig. 4C**). Without ligand, most M_1_R (∼ 75%) occupied the fast Brownian state (*D*_1_ ∼ 0.2 *μm*^2^/*s*), with around 20% in the confined state (*D*_2_ = 0.05–0.10 *μm*^2^/*s*; confinement radius ∼ 160 *nm*) and approximately 5% in the immobile state (*D*_3_ < 0.005 *μm*^2^/*s*) (**Fig. 4D**; **Table 3**; **table S1**). The immobile state had an expected displacement per frame smaller than the localization precision. Ensemble time-averaged mean-squared displacement (ETA-MSD) curves confirmed the free and confined nature of the two mobile states (**fig. S4A,B**).

Confined and immobile fractions were grouped as a non-Brownian population to distinguish them from unhindered Brownian motion. Without ligand, A_2A_R showed higher non-Brownian occupancy (∼ 40%) than M_1_R, with a similar confinement radius (∼ 170 *nm*) (**Fig. 4D,E**; **Table 3**). The confinement size of both receptors were on the scale of membrane organization associated with raft-like nanodomains (36), ordered membrane regions enriched in glycosphingolipids and cholesterol that may serve as receptor–G protein interaction hotspots (22). The C-terminally truncated A_2A_R(1–316)-HaloTag construct had a significantly lower non-Brownian population than full-length Halo-A_2A_R under basal conditions (∼ 23% versus ∼ 37%) (**fig. S6A,B**). Similar effects are reported in hippocampal neurons, where C-terminal truncation reduces A_2A_R confinement and alters its diffusivity, with synapse-associated protein 102 (SAP102) identified as a C-terminal interactor affecting access to compartments of restricted mobility (27).

We compared receptor diffusion after membrane perturbations previously shown to alter membrane order in giant plasma membrane vesicles (37). Methyl-*β*-cyclodextrin (MBCD) decreases the raft-like domain fraction by sequestering membrane cholesterol (38), whereas epigallocatechin gallate (EGCG) increases it by binding lipid headgroups and condensing lipid packing (39). MBCD caused minor changes in M_1_R diffusion but reduced the non-Brownian fraction of A_2A_R toward the basal M_1_R level (**Fig. 4D,E**). EGCG increased non-Brownian occupancy for both receptors relative to their apo states and reduced their diffusion coefficients (**Fig. 4D,E**; **Table 3**). The confinement radii of both M_1_R and A_2A_R decreased to 100– 125 *nm*, consistent with smaller confinement domains after EGCG treatment (**Table 3**).

Carbachol activation increased M_1_R non-Brownian occupancy to around 40%, in the same direction as EGCG, whereas CGS-21680 did not significantly alter A_2A_R diffusion (**Fig. 4D,E**; **Table 3**). One possible explanation is that A_2A_R partitioning into raft-related domains is already near saturation in the basal state, limiting further changes after activation. The higher basal confinement of A_2A_R may contribute to its higher basal activity by increasing local G protein encounter opportunities. Together with receptor conformational dynamics (40), this could facilitate signaling without ligand.

Dual-color single-particle tracking (dcSPT) in cells co-expressing M_1_R–Gα_11_ or A_2A_R–Gα_s_ tested whether receptor confinement increases spatial access to the cognate G protein. Receptor and G protein trajectories showed potential colocalization and interaction regions (**Fig. 5A,B**). G proteins showed different diffusion patterns from their cognate GPCRs, with transitions connecting all three diffusion states (**Fig. 5C**). Their faster Brownian motion was consistent with inner-leaflet anchoring and with the FCS measurements. Despite somewhat slower confined diffusion than their receptor partners, the G proteins had similar confinement radii (∼ 160 *nm*), suggesting access to similarly sized confined membrane regions (**Table 3**).

**Figure 5.**
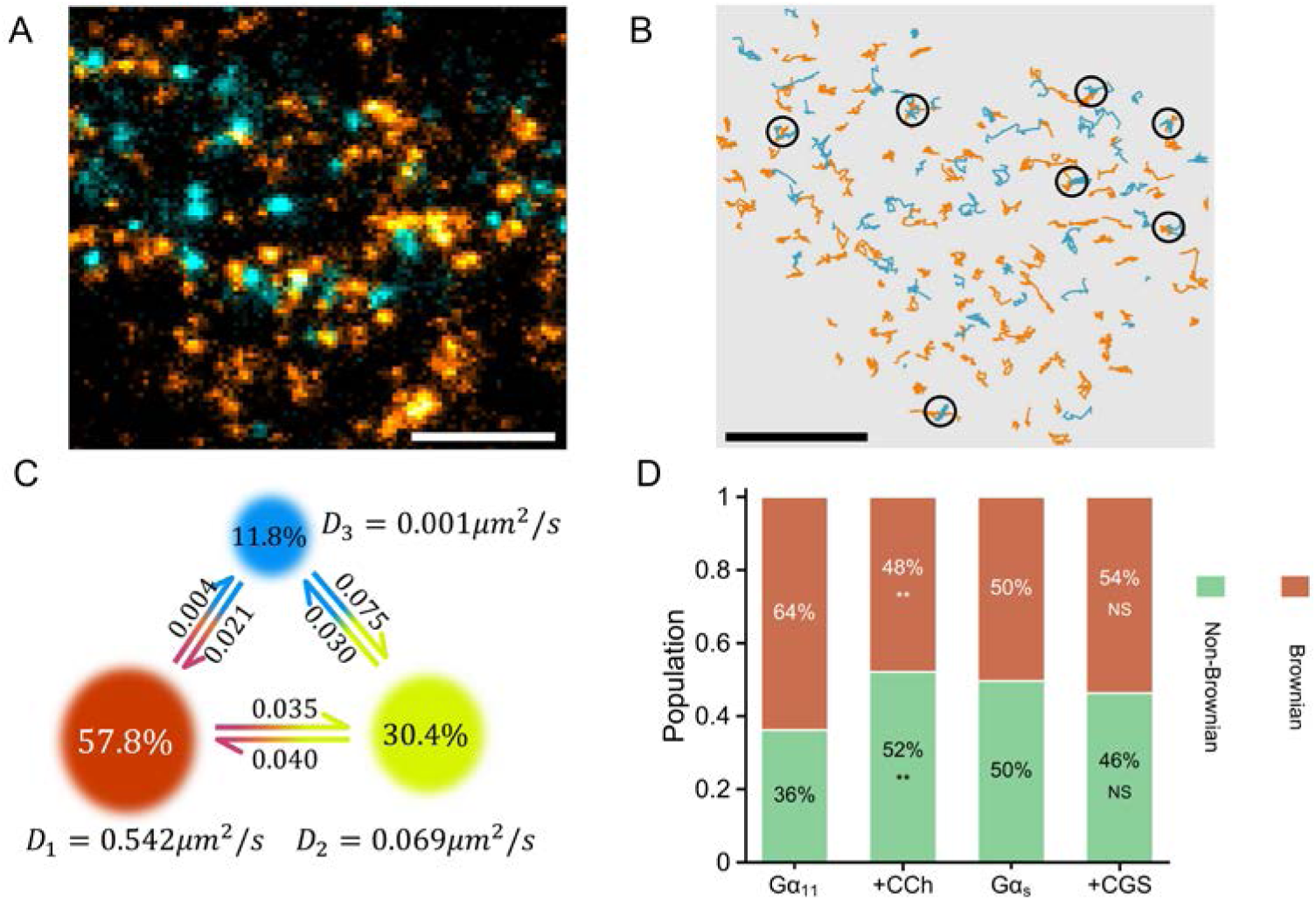
Dual-color tracking of GPCRs and G proteins in live cells. **(A)** Representative frame from a TIRF movie of an apo-state cell co-expressing A_2A_R (orange) and Gα_s_ (cyan). Panels **(A–C)** show this cell, selected from *n* = 18 cells. Scale bar: 5 *μm*. **(B)** Extracted single-molecule trajectories of A_2A_R (orange) and Gα_s_ (cyan), with regions of trajectory overlap marked by dark circles. Scale bar: 5 *μm*. **(C)** G protein trajectories were analyzed using *ExaTrack*, yielding three interchanging diffusion states: *free/Brownian* (*D*_1_ ∼ 0.5 *μm*^2^/*s*), *confined* (*D*_2_ = 0.05–0.10 *μm*^2^/*s*), and *immobile* (*D*_3_ < 0.005 *μm*^2^/*s*). **(D)** Stacked composition bars summarize changes in Brownian (red) and non-Brownian (green) motion of G proteins due to activation of their associated GPCRs by carbachol (CCh) or CGS-21680 (CGS); comparisons are made between agonist treatments and unstimulated conditions (**, *P* < 0.01; NS, not significantly different, tested by two-sided Welch’s *t*-tests with Holm correction). Percentages in **(D)** indicate mean diffusion-population occupancies, with full mean ± SEM values listed in **Table 3** and **table S1**. Flp-In T-REx 293 cells stably expressed Gα_11_-SNAP or Gα_s_-SNAP and transiently expressed Halo-M_1_R or Halo-A_2A_R, respectively. **(D)** *n* = 15 and 13 cells for Gα_11_ without and with carbachol, and *n* = 18 and 13 cells for Gα_s_ without and with CGS-21680, respectively, across three independent biological replicates per condition.

In the apo state, Gα_11_ had a lower non-Brownian fraction (∼ 35%) than Gα_s_ (∼ 50%). Carbachol activation of M_1_R increased Gα_11_ non-Brownian occupancy to approximately 50%, toward the Gα_s_ level, whereas CGS-21680 activation of A_2A_R left Gα_s_ diffusion largely unchanged (**Fig. 5D**; **Table 3**). These trends mirrored those of the receptors (**Fig. 4D,E**), linking confinement with signaling activity. Most observed receptor–G protein colocalizations were transient (0.1–0.5 *s*). Short interaction times and photophysical blinking could both contribute, preventing reliable separation of productive interactions from random colocalization. Rather than assigning specific complex lifetimes (41), we therefore compared time-averaged spatial diffusion maps to assess shared confinement.

Under basal conditions, M_1_R confinement domains were sparse (∼ 25% non-Brownian fraction), whereas Gα_11_ occupied more non-Brownian regions (∼ 35%) in the same cell (**Fig. 6A,B**). White areas in the maps represent regions not traversed by labelled molecules in the recorded trajectories. A_2A_R and Gα_s_ occupied more non-Brownian regions (**Fig. 6C,D**). Corresponding agonist-stimulated maps are shown in **Fig. 6E–H**. Image cross-correlation spectroscopy (ICCS) analysis of the diffusion-state maps yielded spatial auto- and cross-correlation profiles for the receptor and G protein confinement regions (**Fig. 6I**). The effect of the deconvolution correction on the spatial autocorrelation profile is shown in **fig. S5A**. A_2A_R and Gα_s_ showed greater co-confinement (*coco* ≈ 54%) than M_1_R–Gα_11_ (*coco* ≈ 30%) (**Fig. 6J**; **table S2**). This greater spatial overlap could favor productive encounters and contribute to the higher basal signaling of A_2A_R (**Fig. 2B**). The noncognate A_2A_R–Gα_11_ and M_1_R–Gα_s_ control pairs showed low basal co-confinement of approximately 24% and 23%, respectively (**fig. S5B**).

**Figure 6.**
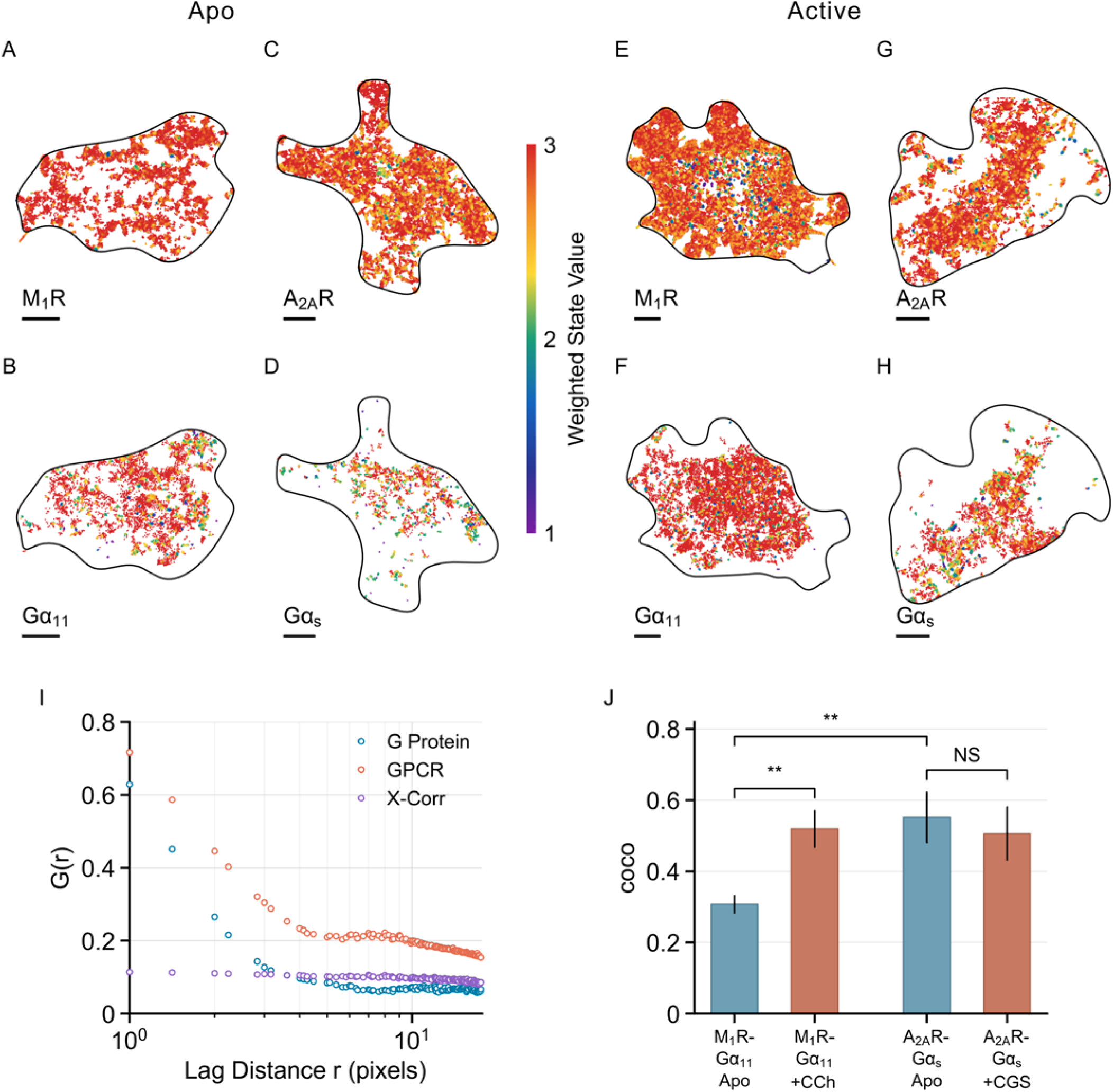
Diffusion-state mapping and GPCR–G protein co-confinement. Representative time-averaged diffusion-state maps generated from dual-color single-particle tracking (dcSPT) trajectories for M_1_R and Gα_11_, or A_2A_R and Gα_s_, in apo and agonist-stimulated states. **(A-D)** Apo (no agonist): M_1_R **(A)**, Gα_11_ **(B)**, A_2A_R **(C)**, Gα_s_ **(D)**; **(E-H)** Active: M_1_R **(E)** and Gα_11_ **(F)** in the presence of 10*μ*M carbachol (+CCh), A_2A_R **(G)** and Gα_s_ **(H)** in the presence of 10*μ*M CGS-21680 (+CGS). Pseudo-color represents the local weighted diffusion-state value *S*′(**r**) (**Eq. 2**) on a 1–3 scale, corresponding to immobile (1), confined (2), and Brownian diffusion (3) states. Cell outlines were obtained from the time averaged images; scale bars, 5 *μ*m. **(I)** Representative ICCS analysis of diffusion maps from one apo A_2A_R–Gα_s_ cell in the non-Brownian diffusion regions (*S*′(**r**) < 2.25): autocorrelations of receptors (orange) and G proteins (blue) and their cross-correlation (X-corr, violet). **(J)** Co-confinement fractions (*coco*) for M_1_R–Gα_11_ and A_2A_R–Gα_s_ in apo and active states, calculated from the corrected ICCS amplitudes (**Eq. 3**). Data are mean ± SEM; *coco* values are listed in **table S2**. **, *P* < 0.01; NS, not significant, tested by two-sided Welch’s *t*-tests with Holm correction. Flp-In T-REx 293 cells stably expressed the SNAP-tagged G protein and transiently expressed the cognate Halo-GPCR. Panels **(A,B)**, **(C,D,I)**, **(E,F)**, and **(G,H)** represent datasets of *n* = 15, 18, 13, and 13 cells, respectively; these counts also apply to the corresponding conditions in **(J)**. Each condition was collected across three independent biological replicates.

Carbachol stimulation increased M_1_R confinement and produced Gα_11_ confinement in similar membrane regions, increasing their co-confinement to approximately 50% (**Fig. 6E,F,J**; **table S2**). A_2A_R and Gα_s_ were already extensively co-confined under basal conditions, and CGS-21680 did not significantly change their overlap (*coco* ≈ 48%) (**Fig. 6G,H,J**). These qualitative patterns persisted in G-protein-depleted HEK293 cell lines: A_2A_R–Gα_s_ remained highly co-confined in *ΔG_s_* cells with little agonist-dependent change, whereas carbachol increased M_1_R–Gα_11_ co-confinement in *ΔG_q_*_/11_ cells (**fig. S5B**; **tables S2 and S4**). At the lower expression levels used for dcSPT, M_1_R–Gα_11_ co-confinement showed no significant association with apparent receptor or G-protein density over the measured range (**fig. S5C,D**). Together, these findings link basal and agonist-induced signaling to shared receptor–G protein confinement within membrane nanodomains.

### Linking GPCR confinement to constitutive activity

The co-confinement measurements suggest that membrane organization affects receptor access to G proteins but do not establish whether changing that organization alters signaling. In Halo-A_2A_R-expressing cells without orthosteric ligand, EGCG increased basal cAMP production over the concentration range tested, whereas MBCD reduced it (**Fig. 7A**). Although these agents broadly perturb the membrane, the agreement between their effects on confinement and cAMP production indicates that constitutive A_2A_R signaling is sensitive to membrane organization.

**Figure 7.**
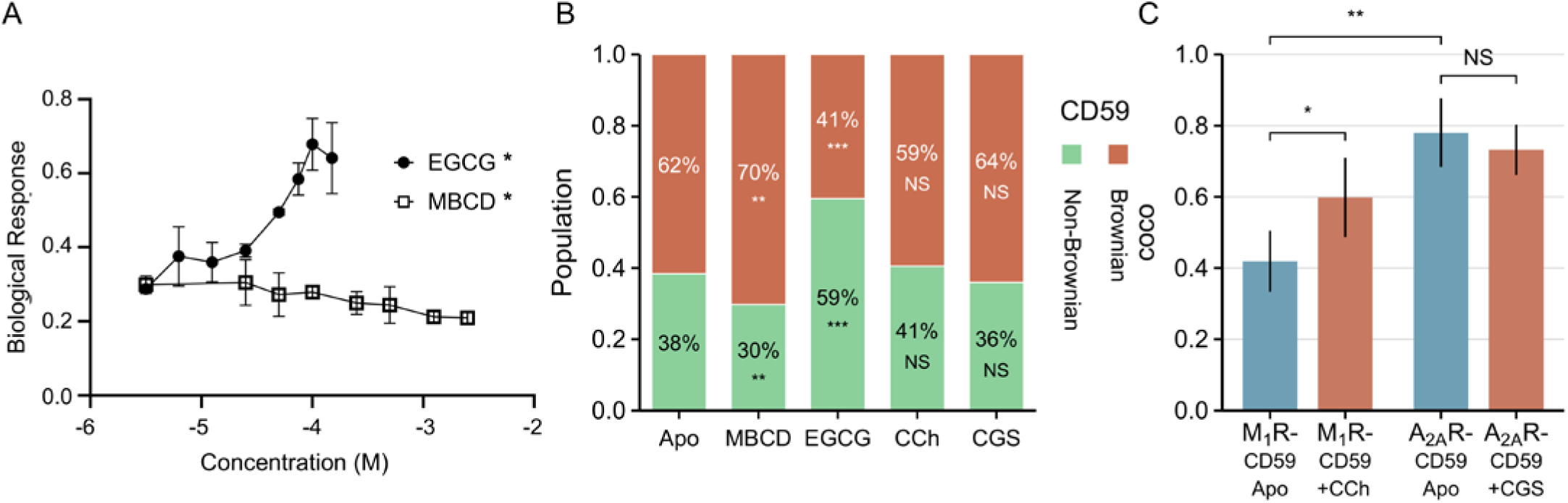
Functional effects of membrane perturbation and CD59-based characterization of GPCR membrane organization. **(A)** A_2A_R-mediated cAMP signaling measured after treatment of Flp-In T-REx 293 cells stably expressing Halo-A_2A_R with EGCG or MBCD at the indicated concentrations. Data are mean ± standard deviation (SD) from *n* = 3 independent biological experiments per condition/concentration and normalized between 0 and 1, as defined by saturating inverse agonist and agonist conditions, respectively (Fig. 2B). *E*_max_ values were compared between each treatment and its vehicle control using two-sided Welch’s *t*-tests. **(B)** Diffusion populations in Flp-In T-REx 293 cells stably expressing Halo-CD59 under Apo, MBCD, EGCG, carbachol (+CCh), and CGS-21680 (+CGS) conditions, determined by SPT analysis. Stacked bars show mean Brownian (red) and non-Brownian (green) populations. Detailed diffusion parameters, including means ± SEM, are listed in **Table 4** and **table S3**. **(C)** Co-confinement fractions (*coco*) of M_1_R or A_2A_R with CD59 in apo and agonist-stimulated conditions, determined by ICCS; *coco* values are listed in **table S2**. *, *P* < 0.05; **, *P* < 0.01; ***, *P* < 0.005; NS, not significant, tested by two-sided Welch’s *t*-tests with Holm correction. **(B)** *n* = 14, 17, 14, 11, and 8 cells for Apo, MBCD, EGCG, carbachol, and CGS-21680, respectively. **(C)** Data are mean ± SEM from stable Halo-CD59 Flp-In T-REx 293 cells transiently expressing SNAP-M_1_R or SNAP-A_2A_R; *n* = 9 and 11 cells for M_1_R–CD59 without and with carbachol, and *n* = 8 cells each for A_2A_R–CD59 without and with CGS-21680. Each condition in **(B,C)** was collected across three independent biological replicates.

**Table 4.** SPT analysis for CD59 under different conditions^a^.

| **Protein** | **Condition** | $D_{Br}$ ($\mu\text{m}^2/\text{s}$) | $D_{conf}$ ($\mu\text{m}^2/\text{s}$) | $f_{NBr}$ (%)<sup>b</sup> | $R_{conf}$ (nm)<sup>c</sup> | **n** (cells) |
| --- | --- | --- | --- | --- | --- | --- |
| CD59 | Apo | $0.224 \pm 0.010$ | $0.066 \pm 0.037$ | $38.4 \pm 7.5$ | $150 \pm 7$ | 14 |
| | MBCD | $0.231 \pm 0.007$ | $0.075 \pm 0.014$ | $29.8 \pm 3.3$ | $232 \pm 16$ | 17 |
| | EGCG | $0.197 \pm 0.016$ | $0.065 \pm 0.019$ | $59.5 \pm 8.5$ | $137 \pm 5$ | 14 |
| | Carbachol | $0.228 \pm 0.013$ | $0.086 \pm 0.024$ | $40.5 \pm 10.6$ | $176 \pm 11$ | 11 |
| | CGS-21680 | $0.239 \pm 0.005$ | $0.083 \pm 0.024$ | $36.0 \pm 6.9$ | $150 \pm 12$ | 8 |

<sup>a</sup> SPT data was analyzed using ExaTrack (35); for each condition, the weighted mean values across cells are listed, with errors reported as ± SEM. <sup>b</sup> $f_{NBr}$ , percentage of the non-Brownian population, calculated as the sum of the confined and immobile fractions. <sup>c</sup> $R_{conf}$ , confinement radius of the confined diffusion population, calculated using **Eq. S14**

To examine the nature of these confinement regions, we studied the raft-associated, GPI-anchored outer-leaflet protein CD59 in Halo-CD59-expressing cells. In the apo state, approximately 40% of CD59 molecules occupied non-Brownian states, with a confinement radius of around 150 *nm*. MBCD reduced the non-Brownian fraction to around 30% and increased its confinement radius to about 230 *nm*. EGCG instead increased the non-Brownian fraction to approximately 60%, with a smaller confinement radius of around 140 *nm* (**Fig. 7B**; **Table 4**; **table S3**). These opposing responses support CD59 as an independent marker of raft-related nanodomain organization. Carbachol and CGS-21680 did not significantly alter CD59 diffusion, indicating that agonist-dependent receptor changes were not accompanied by global reorganization of CD59-marked nanodomains.

In cells co-expressing Halo-CD59 with SNAP-M_1_R or SNAP-A_2A_R, basal M_1_R–CD59 co-confinement was approximately 40%, whereas A_2A_R–CD59 co-confinement was approximately 80%. Carbachol increased M_1_R–CD59 co-confinement to around 60%, whereas A_2A_R–CD59 co-confinement remained high after CGS-21680 treatment at approximately 75% (**Fig. 7C**; **table S2**). These receptor-dependent patterns matched those observed with their cognate G proteins. Spatial correspondence with CD59 and matched responses to MBCD and EGCG support the association of receptor confinement with raft-related membrane organization.

## DISCUSSION

Basal activity has been reported for many GPCRs (4, 6, 7, 10, 12, 13, 42), but the molecular mechanism is often attributed primarily to the structural plasticity of the receptor, allowing spontaneous transitions to active conformations (43, 44). A_2A_R studies using ^19^F nuclear magnetic resonance (NMR) and single-molecule FRET show that this receptor exchange between inactive and active conformations on microsecond-to-millisecond timescales, providing a plausible explanation for basal activity (40, 45, 46). At the same time, work on C-terminally truncated A_2A_R shows that minor changes in receptor architecture can diminish or even abolish basal cAMP production in live cells, despite ligand binding and G protein coupling in vitro being largely preserved (47). For muscarinic receptors, NMR studies depict a similarly complex energy landscape for the M_2_ muscarinic receptor (M_2_R) (48), but apo/antagonist-bound receptors occupy active states to a smaller extent than A_2A_R. FRET-based M_1_R biosensors indicate robust agonist-induced activation yet low basal activation signals, consistent with an ensemble biased toward inactive conformations (49, 50).

This study adds a spatiotemporal dimension by directly linking basal signaling of A_2A_R and M_1_R to real-time receptor–G protein interactions and nanoscale membrane organization. Previous single-particle tracking work on A_2A_R in neurons identifies a two-state diffusion model in which agonist stimulation promotes receptor transition into compartments characterized by slow diffusion, access to which depends on cholesterol, the C-terminus, and associated scaffolds (27). These confined regions are interpreted as cholesterol-rich microdomains compatible with lipid rafts (26). More generally, experimental and computational studies of liquid-ordered nanodomains indicate that proteins within these regions diffuse more slowly and are locally enriched, increasing the frequency of collisions among signaling components (51, 52). Studies of GPCR–lipid interactions and membrane-mediated oligomerization further emphasize that nanodomains and local lipid composition can strongly alter GPCR conformation, oligomerization, and signaling efficiency (53, 54).

Within this framework, our FCCS and SPT measurements suggest that differences in basal activity among GPCRs arise from the dynamic organization of receptors and G proteins within the plasma membrane. Cross-correlation data at multiple sites in several cells show that, in the basal state, only a minor fraction of M_1_R co-diffuses with Gα_11_. In contrast, the majority of A_2A_R co-diffuse with Gα_s_ in the absence of ligands. As expression levels are generally high and often different between GPCRs and their cognate G proteins in these experiments, the co-diffusion fractions should be taken as upper limits for the fractions of (pre)coupled RG complexes. In addition, FCCS does not unambiguously distinguish true co-diffusion from co-confinement of receptors and G proteins within shared membrane nanodomains on the scale of the confocal volume (∼ 300 nm). The differences between the co-diffusing and slow-diffusing fractions of both Gα_11_ and Gα_s_ also suggest the involvement of transient interactions (**Table 2**). Supporting this, FCCS measurements of A_2A_R and Gα_s_ in the apo state within dimmer cellular regions revealed a lower fraction of RG complexes (**fig. S3A,B**). In contrast, at the lower expression levels used for dual-color SPT, M_1_R–Gα_11_ co-confinement showed no detectable correlation with either receptor or G protein density over the measured range (**fig. S5C,D**), indicating that the observed co-confinement was not primarily determined by expression level. The present analysis focuses primarily on the canonical M_1_R–Gα_11_ and A_2A_R–Gα_s_ pairs. The lower co-confinement observed for the noncognate A_2A_R–Gα_11_ and M_1_R–Gα_s_ pairs, together with preservation of the cognate co-confinement patterns in *ΔG_s_* and *ΔG_q_*_/11_ cells (**fig. S5B**), argues against these trends arising predominantly from nonspecific crosstalk or competition with endogenous cognate G proteins. Other endogenous G-protein interactions, however, cannot be excluded.

Dual-color particle tracking further refines this model by elucidating how receptors and G proteins distribute among distinct diffusion states across the plasma membrane. Both receptors and G proteins show a fast Brownian state, an intermediate confined state, and an immobile state. In the absence of ligands, A_2A_R and Gα_s_ display higher non-Brownian fractions than M_1_R and Gα_11_, with comparable confinement radii of ∼ 160 nm. This confinement scale is consistent with, but does not by itself establish, partitioning into cholesterol- andF sphingolipid-enriched liquid-ordered nanodomains (20). The measured *R*_conf_ represents an effective spatial extent of restricted motion rather than the physical radius of an individual nanodomain. Its magnitude may also be broadened by the finite temporal resolution and motion blur of the measurement. The observed confinement could reflect multiple smaller ordered domains within a larger membrane compartment, cortical actin-defined corrals, or a combination of these mechanisms. Receptor-specific structural features also contribute to this behavior: the C-terminally truncated A_2A_R(1–316)-HaloTag construct showed a lower non-Brownian population than full-length A_2A_R under basal conditions (**fig. S6B**), consistent with previous evidence that the A_2A_R C-terminus and associated protein interactions influence receptor access to slowly diffusing membrane compartments (27). Cholesterol depletion with MBCD reduces the non-Brownian fraction of A_2A_R while exerting only minor effects on M_1_R. In contrast, targeting lipid rafts with EGCG increases the non-Brownian fractions of M_1_R and A_2A_R to approximately 50% and 70%, respectively, and reduces their confinement radii to approximately 100–125 nm (**Table 3**), consistent with increased coverage of smaller ordered domains. Together, these observations indicate that GPCR confinement reflects contributions from both membrane lipid organization and receptor-specific molecular interactions.

The CD59 measurements provide an independent physical reference for this interpretation (**Fig. 7B,C**). CD59 and A_2A_R showed similar basal confinement and responded similarly to MBCD and to EGCG. A_2A_R exhibited much higher basal co-confinement with CD59 than M_1_R; agonist activation increased M_1_R–CD59 co-confinement whereas A_2A_R remained high, paralleling the receptor–G-protein co-confinement trends. Because CD59 is anchored to the outer membrane leaflet, whereas G proteins are associated with the cytoplasmic leaflet, this correspondence also suggests membrane organization is coupled across the bilayer (21, 23). The opposing effects of EGCG and MBCD on basal A_2A_R-mediated cAMP production matched the direction of their effects on receptor confinement, supporting a link between raft-related membrane organization and constitutive signaling. Although these agents act broadly on the plasma membrane, the agreement between the tracking and functional measurements shows that nanoscale membrane organization can alter constitutive A_2A_R activity.

Agonist binding can alter the exposure of palmitoylated and intracellular receptor regions, increasing affinity for cholesterol-/sphingolipid-rich and anionic membrane nanodomains and promoting recruitment of receptor–G-protein complexes (55, 56). The coordinated increases in M_1_R confinement and its co-confinement with both CD59 and Gα_11_ are consistent with activation-dependent redistribution into pre-existing membrane nanodomains. In contrast, A_2A_R already occupied these regions substantially in the basal state and changed little after agonist treatment. The absence of agonist-dependent changes in CD59 diffusion further supports receptor redistribution rather than global reorganization of the nanodomains themselves. Their partial correspondence with CD59 suggests that raft-related membrane organization is a major, but not exclusive, contributor to GPCR confinement. Cortical actin meshes, protein scaffolds, receptor assemblies, and *β*-arrestin-dependent sorting into clathrin-coated pits may contribute to the remaining confined regions (17, 18, 57).

Following GPCR activation, both G proteins also show an increased average confinement radius of approximately 200–250 nm. One speculative possibility is that depalmitoylation and dissociation of *Gα* and *Gβγ* from the receptor allow the activated subunits to sample adjacent membrane corrals and downstream effectors. Alternatively, competition with *β*-arrestin/clathrin for activated receptors may shorten the G-protein residence time near receptor-containing regions and release it into other membrane compartments. Future experiments involving simultaneous tracking of *β*-arrestin will help distinguish these possibilities.

The effective encounter rate *k_enc_* between GPCRs and G proteins is determined not only by their overall expression levels, but also by the interplay of local surface density and diffusion within and outside liquid-ordered nanodomains. In less ordered membrane regions, receptors and G proteins diffuse rapidly but are more sparsely distributed, limiting the time during which they remain close enough to interact. Shared confinement of the receptor and its G protein within raft-related membrane regions locally enriches both species while restricting their movement to a smaller membrane area. This confinement increases their residence time within a shared region and enhances the probability of repeated and potentially productive encounters, despite the slower diffusion. When the Brownian and confined-state diffusion parameters from **Table 3** are applied to the encounter-rate model, the estimated rate increases from 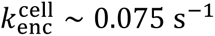 across the cell–substrate contact region to 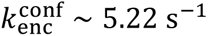 within confined membrane domains. The corresponding mean encounter time decreases from approximately 13.3 s to 0.19 s, providing a quantitative link between nanoscale membrane organization, receptor–G-protein encounters, and constitutive signaling.

We propose a two-factor model in which constitutive GPCR activity depends on both (*i*) adequate conformational flexibility to populate signaling-competent states and (*ii*) sufficient partitioning of the receptor and its G protein into liquid-ordered membrane nanodomains, which increases the probability of local receptor–G-protein encounters. A_2A_R fulfills both criteria under basal conditions, maintaining a substantial population of active conformers and exhibiting robust co-confinement with Gα_s_ within these nanodomains. This leads to high encounter rates and substantial constitutive cAMP production. In contrast, the Halo-M_1_R-expressing system showed very low basal Gα_q/11_–Ca^2+^ signaling under our conditions, consistent with lower basal occupancy of active receptor conformers and limited M_1_R–Gα_11_ co-confinement; both increased upon agonist binding. Future high-resolution conformational studies on full-length M_1_R, using approaches such as NMR or single-molecule FRET, will be needed to test this model and quantify its basal conformational ensemble relative to A_2A_R.

The two-factor model implies that GPCR activity in cells is determined not only by the intrinsic conformational properties of the receptor, but also by the spatiotemporal organization of the receptor and its signaling effectors within the membrane. This framework may help explain why the same receptor displays signaling bias and promiscuity in different cellular contexts (44, 58). It also suggests that combining ligands that alter receptor conformation with approaches that alter liquid-ordered nanodomain organization or receptor partitioning may enable more precise adjustment of tonic signaling and enhance therapeutic selectivity.

## MATERIALS AND METHODS

### Plasmid constructs

The Halo-M_1_R plasmid was constructed as described previously (59). Complementary DNAs (cDNAs) for Halo-A_2A_R, the C-terminally truncated A_2A_R(1–316)-HaloTag construct, Halo-CD59, Gα_11_-SNAP, and Gα_s_-SNAP were synthesized by GenScript after human codon optimization. The A_2A_R constructs contained an N-terminal metabotropic glutamate receptor 5 (mGluR5)-derived signal peptide (MVLLLILSVLLLKEDVRGSAQS), hemagglutinin (HA) tag, CD8A-derived sequence (MALPVTALLLPLIALLLHAAR), and Myc tag upstream of the receptor sequence. Full-length Halo-A_2A_R contained A_2A_R residues M1–S412, whereas the truncated construct contained residues M1–A316 followed by a C-terminal HaloTag. The Halo-CD59 coding sequence was subcloned into the pcDNA^TM^5/FRT/TO mammalian expression vector. For Gα_11_-SNAP and Gα_s_-SNAP, SNAPf-Tag was inserted in the *αA*–*αB* loop of the *α*-helical domain between positions 97 and 98 of Gα_11_, and positions 113 and 114 of the Gα_s_ long isoform, respectively (60). The G protein inserts were excised with NheI and NotI and subcloned into pcDNA^TM^5/FRT/TO.

To create SNAP-M_1_R and SNAP-A_2A_R, the HaloTag sequence was replaced with the SNAPf-tag sequence from Gα_s_-SNAP using NEBuilder HiFi DNA Assembly Master Mix (New England Biolabs [NEB], E2621S). Primers were designed using NEBuilder Assembly Tool v2.11.2 (https://nebuilder.neb.com/). Halo-M_1_R and Halo-A_2A_R were linearized by polymerase chain reaction (PCR) using Q5 High-Fidelity 2X Master Mix (NEB, M0492S) with builder-M1-fwd 5’-cctgcaggtataggcgcg-3’ and builder-M1-rev 5’- atcgatgccggccaggtc-3’ for M_1_R, and builder-A2A-fwd 5’-cctgccggaattggagcc-3’ and builder-A2A-rev 5’- atcgattccggccaggtc-3’ for A_2A_R. PCR products were treated with DpnI (NEB, R0176S) and purified by gel extraction (NEB, T1120S). The SNAPf-tag sequence was amplified with SNAP-M1-fwd 5’- aggacctggccggcatcgatatggacaagg actgcgagatgaaaag-3’ and SNAP-M1-rev 5’- ggcgcgcctatacctgcagggcccaggcca ggtttgcc-3’ for M_1_R, and SNAP-A2A-fwd 5’- aggacctggccggaatcgatatggacaagg actgcgagatgaaaag-3’ and SNAP-A2A-rev 5’- ggggctccaattccggcagggcccaggcca ggtttgcc-3’ for A_2A_R, followed by gel extraction. NEBuilder reactions were assembled according to the manufacturer’s instructions, transformed into chemically competent *Escherichia coli* Top10 cells, and plated on LB Lennox agar (Bioshop, LBL405) containing 100 *μ*g/mL carbenicillin (Fisher BioReagents, BP2648). All cloning steps were confirmed by sequencing.

### Cell culture and transfection

Most experiments were performed in Flp-In T-REx 293 cells (Invitrogen, R78007), cultured and selected as described previously (59). Stable cell lines expressing Halo-M_1_R, Halo-A_2A_R, Gα_11_-SNAP, Gα_s_-SNAP, or Halo-CD59 were generated using the Flp-In T-REx system. Cells were seeded in 6-cm dishes at 1 × 10^6^ cells per dish and transfected at ∼ 70% confluency with pcDNA5/FRT/TO carrying the protein of interest and pOG44 (Invitrogen, V600520) in a 1:9 ratio using 12 *μ*L Lipofectamine 2000 (Invitrogen, 11668027). Hygromycin-resistant colonies were pooled after ∼ 10–14 days of selection.

Stable Halo-M_1_R and Halo-A_2A_R cells were used for the live-cell signaling assays and for single-color fluorescence measurements. Stable Gα_11_-SNAP and Gα_s_-SNAP cells were used for single-color G protein measurements, and stable Halo-CD59 cells were used for single-color CD59 measurements. For fluorescence experiments, cells were seeded on ultraclean 35-mm glass-bottom *μ*-dishes (Ibidi, 81158) at ∼ 10^5^ cells per dish, grown to 50–70% confluency, and induced with 1–100 ng/mL doxycycline (Sigma-Aldrich, PHR1145) for 8 h.

For the main dual-color GPCR–G protein experiments, stable Gα_11_-SNAP or Gα_s_-SNAP Flp-In T-REx 293 cells were transiently transfected with Halo-M_1_R or Halo-A_2A_R, respectively. For dual-color GPCR–CD59 experiments, stable Halo-CD59 cells were transiently transfected with SNAP-M_1_R or SNAP-A_2A_R. After 12 h of adhesion, cells were transfected with 1 *μ*g of the appropriate GPCR plasmid using 4 *μ*L Lipofectamine 2000 per dish. Six hours after transfection, the medium was replaced with minimal growth medium supplemented with 100 ng/mL doxycycline, and cells were labelled and imaged 12 h later.

The *ΔG_s_* and *ΔG_q_*_/11_ HEK293 cell lines (61, 62) were used only for the supplementary GPCR–G protein control experiments indicated in **fig. S5B–D** and **table S4**. In these cell lines, both the relevant Halo-tagged GPCR and SNAP-tagged G protein were introduced by transient cotransfection. For the supplementary A_2A_R C-terminal truncation experiment, Halo-A_2A_R was transiently transfected into Flp-In T-REx 293 cells under the same conditions described above.

### Signaling assays

For calcium flux assays, stably transfected Halo-M_1_R cells were seeded in poly-D-lysine-coated glass-bottom 96-well plates with black walls (Greiner, 655090) at ∼ 75,000 cells per well. After overnight adhesion, cells were induced with 1 *μ*g/mL doxycycline for 24 h, loaded with 3 *μ*M Fura-2 AM Ca^2+^ dye (Invitrogen, F1201) in Hank’s Balanced Salt Solution (HBSS, Cytiva, SH30268.01) for 45 min at 37 ^∘^C in the dark, washed twice with HBSS, and incubated for another 15 min at 20 ^∘^C for de-esterification. The inverse agonist pirenzepine (Thermo Scientific Chemicals, J62252-MC) was included during de-esterification, while the agonist carbachol (MilliporeSigma, 21238) was added immediately before measurement. Fura-2 emission at 510 nm was measured on a Cytation S63 spectrophotometer (BioTek) after excitation at 340 and 380 nm, and calcium responses were calculated from the 340/380 intensity ratio (63).

cAMP signaling assays were performed according to the cAMP-Glo Max protocol (Promega, V1681). Stably transfected Halo-A_2A_R cells were seeded in poly-D-lysine-coated glass-bottom 96-well plates with white walls (Thermo Scientific, 165306) at ∼ 10,000 cells per well and induced for 24 h with 1 *μ*g/mL doxycycline in the presence of 100 ng/mL pertussis toxin (PTX, Invitrogen, PHZ1174) to inactivate endogenous Gα_i_ and Gα_o_ proteins. Cells were treated for 30 min with the agonist CGS-21680 (MilliporeSigma, C141) or the inverse agonist ZM-241385 (MilliporeSigma, Z0153) in Dulbecco’s phosphate-buffered saline (D-PBS; Gibco, 14190144) containing cAMP phosphodiesterase inhibitors. cAMP-Glo Max lysis/detection reagent and Kinase-Glo reagent were then added sequentially for 20 min each. Luminescence was measured on the Cytation S63 spectrophotometer. Vehicle-control wells received an equivalent volume of dimethyl sulfoxide (DMSO), matching the final DMSO concentration in the corresponding treatment wells. Both assays were performed in biological triplicate, and concentration-response curves were fitted in GraphPad Prism 10 (GraphPad, CA) to estimate *EC*_50_ values.

### Fluorescent labelling and ligand treatment

Cells expressing Halo-M_1_R or Halo-A_2A_R were labelled with 1 nM JF549i-HTL (HaloTag ligand) in complete culture medium, following the previously described HaloTag labelling workflow (59). SNAP-M_1_R and SNAP-A_2A_R were labelled with 50 nM JF549i-STL (SNAP-tag ligand) in complete culture medium for 30 min at 37 ^°^C. Gα_11_-SNAP and Gα_s_-SNAP were labelled with 50 nM membrane-permeable JF669-STL in complete culture medium for 30 min at 37 ^°^C. Halo-CD59 was labelled with 1 nM JF635i-HTL in complete culture medium for 5 min at 37 ^°^C. For the C-terminal truncation measurements, A_2A_R(1–316)-HaloTag was labelled with 1 nM JF635-HTL under the same conditions. All fluorophores were obtained from Janelia Farm.

Cells were washed several times at 15-min intervals to remove free dye and transferred to 1 mL FluoroBrite Dulbecco’s modified Eagle’s medium (DMEM) for imaging. For dual-color GPCR–G protein experiments, SNAP-tag labelling of G proteins was performed before HaloTag labelling of GPCRs.

For receptor activation, cells expressing M_1_R were incubated with 10 *μ*M carbachol, and cells expressing A_2A_R were incubated with 10 *μ*M CGS-21680. Both agonist treatments were performed in complete culture medium for 30 min at 37 ^∘^C. To perturb plasma membrane organization, live cells were incubated with 50 *μ*M EGCG (Cayman Chemical, 0531242-85) or MBCD (MilliporeSigma, C4555) in complete culture medium for 2 h at 37 ^∘^C.

### Fluorescence Correlation Spectroscopy

FCS measurements in live cells were performed on a custom-built confocal microscope equipped with a hardware correlator, as described previously (31). Single-color FCS measurements were performed using the corresponding stable GPCR or G protein cell lines. Dual-color FCS measurements were performed in stable Gα_11_-SNAP or Gα_s_-SNAP cells transiently expressing Halo-M_1_R or Halo-A_2A_R, respectively.

Before collecting correlation data, a confocal scan was used to identify cells exhibiting fluorescence signals of 10^4^–10^5^ photons/s at excitation intensities around 100 W⋅cm^−2^. For dual-color measurements, the input powers of the green (532 nm) and red (638 nm) lasers were adjusted so that the two emission signals had similar intensities. Within a 50 *μ*m × 50 *μ*m field of view, multiple regions on the basal membrane were selected at protein surface densities of approximately 1–100 molecules/*μ*m^2^; the mean receptor and G-protein densities for each dual-color FCCS condition are reported in Table 2. Correlation curves were acquired for 20 s, with at least 10 repeats at each location. Correlation curves from single- and dual-color FCS were analyzed using a custom-written MATLAB program (31). Calibration procedures, fitting models, parameter definitions, and equations are provided in Supplementary Materials and Methods (**Eqs. S1 to S7**).

### Single-Particle Tracking

Single-molecule imaging of GPCRs, G proteins, and CD59 in live cells was performed on a custom-built total internal reflection fluorescence (TIRF) microscope described previously (59). The stable and transiently transfected cell preparations used for single- and dual-color measurements were as described above. For TIRF experiments, 35-mm dishes containing fluorescently labelled cells were mounted on a custom-built sample holder. Only adherent, non-mitotic cells with intact morphology and a sufficiently flat basal membrane were selected for imaging. Rounded or dividing cells, highly confluent regions, cells with pronounced membrane folds, and regions containing large intracellular vesicles were excluded from analysis.

GPCRs labelled with JF549i-HTL, JF549i-STL or JF635-HTL and CD59 labelled with JF635i-HTL were imaged at an excitation intensity of 150 W/cm^2^. G proteins labelled with JF669-STL were imaged at 750 W/cm^2^ to obtain a comparable signal-to-noise ratio (SNR ∼ 10). These conditions corresponded to a localization precision of approximately 25 nm and a diffusion detection limit of *D* > 0.005 *μ*m^2^/s. Live-cell TIRF movies were acquired for 1–2 min at 30 ms per frame (33 fps).

Individual fluorescent particles were detected and tracked in time and space using the TrackMate Linear Assignment Problem algorithm implemented in Fiji (64), as described previously (59). Particles were tracked until they photobleached, disappeared, or merged with neighbouring particles. Trajectories containing at least five steps were analyzed with *ExaTrack*. Building on *ExTrack* (65), *ExaTrack* models Brownian, directed, and confined motion, transitions between motion states, and localization error (35). In the confined-motion model, the tracked particle moves within a quadratic potential well whose center can itself diffuse. Particle-linking parameters, model selection, confinement analysis, and ensemble time-averaged mean-squared displacement validation are described in Supplementary Materials and Methods. Statistical comparisons of SPT-derived diffusion-state populations were performed at the cell level using two-sided Welch’s *t*-tests, with Holm correction for multiple comparisons.

Time-averaged spatial maps of diffusion states were generated from the posterior state probabilities returned by ExaTrack. For localization *t* in trajectory *k*, the continuous probability-weighted diffusion-state index was calculated as

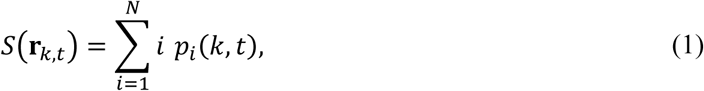

where *p_i_*(*k*, *t*) is the posterior probability of state *i*, as defined in **Eq. 1**. To reduce local sampling noise and highlight mesoscale spatial organization, neighbouring localizations within a one-pixel radius (0.167 *μ*m) were identified, and the smoothed diffusion-state index was calculated as

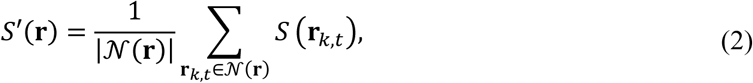

as defined in **Eq. 2**. Diffusion maps were displayed on a 1–3 scale corresponding to immobile, confined, and free/Brownian diffusion, respectively. Cell boundaries obtained from the corresponding time-averaged TIRF images were overlaid as black contours.

### Image Cross-Correlation Spectroscopy

ICCS was applied to paired diffusion-state maps to quantify spatial co-confinement between GPCRs and G proteins or between GPCRs and CD59. The input for each spectral channel was the smoothed diffusion-state map *S*′(**r**) defined in **Eq. 2**, restricted to the common cell contour. Positions with *S*′(**r**) < 2.25 were retained to emphasize immobile and confined regions.

For each channel, mean-subtracted diffusion-state fluctuations *δM_a_*(**r**) and normalized spatial auto- and cross-covariance functions *g_ab_*(*Δ***r**) were calculated. Because the diffusion maps were time averaged, the correlations were evaluated only as functions of spatial displacement. Radial averaging yielded the one-dimensional spatial correlation profiles *G_ab_*(*r*). The measured autocorrelation profiles were modelled as point-spread-function convolutions and corrected by Wiener–Tikhonov deconvolution using a simulated point-spread-function baseline.

The corrected receptor autocorrelation amplitude, *g_R_*(0), and the receptor–partner cross-correlation amplitude, *g_x_*(0), were estimated from correlation values at radial lags *r* < 10 pixels. The co-confinement fraction was calculated as

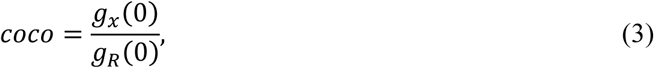

as defined in **Eq. 3**. Detailed map filtering, covariance, radial averaging, deconvolution, and amplitude calculations are provided in Supplementary Materials and Methods. Statistical comparisons of cell-level co-confinement fractions were performed using two-sided Welch’s *t*-tests, with Holm correction for multiple comparisons.

Mean encounter rates were estimated from the measured diffusion coefficients and local G protein densities using two-dimensional Smoluchowski theory, as detailed in Supplementary Materials and Methods. Custom code used for FCS/FCCS fitting and analysis, ICCS analysis, diffusion-map generation, and downstream diffusion analyses is archived at Zenodo (https://doi.org/10.5281/zenodo.23148231).

## SUPPLEMENTARY MATERIALS

Materials and Methods

Figs. S1 to S6

Tables S1 to S4

References (66) to (70)

## ACKNOWLEDGMENTS

We thank Dr. Balmiki Kumar from the Department of Chemistry & Physical Sciences at University of Toronto for assistance with live cell signaling assays, Dr. Luke Lavis from Janelia Research Campus for gifting us the JF549i-HTL, JF635-HTL, JF635i-HTL and JF669-STL fluorophores, and Dr. Asuka Inoue and the Inoue laboratory at Tohoku University for providing the *ΔG_s_* and *ΔG_q_*_/11_ HEK293 cell lines.

## Author contributions

Conceptualization: XZ, CCG; Methodology: XZ, MS, FS, AB, LEW, CCG; Investigation: XZ, LP, GD, MLU; Visualization: XZ, MS, AB; Funding acquisition: CCG, RSP, LEW; Project administration: XZ, CCG; Supervision: CCG, RSP, LEW; Writing – original draft: XZ, CCG; Writing – review & editing: XZ, MS, LP, FS, AB, MLU, LEW, RSP, CCG.

## Data, code and materials availability

Custom code for FCS/FCCS fitting and analysis, ICCS analysis, diffusion-map generation, and downstream diffusion analyses is publicly available on Zenodo (https://doi.org/10.5281/zenodo.23148231). The archive includes the ExaTrack source used in this study, with credit to its original developer, François Simon. Complete sequences of nine plasmid constructs have been submitted to GenBank under submission ID SUB16534627; accession assignment and public release are pending. All other data needed to evaluate the conclusions in the paper are present in the paper or the Supplementary Materials.

## REFERENCES

1. R. Santos et al., A comprehensive map of molecular drug targets. Nature Reviews Drug Discovery 16, 19–34 (2017).

2. D. M. Rosenbaum, S. G. F. Rasmussen, B. K. Kobilka, The structure and function of G-protein-coupled receptors. Nature 459, 356–363 (2009).

3. G. Milligan, Constitutive Activity and Inverse Agonists of G Protein-Coupled Receptors: a Current Perspective. Molecular Pharmacology 64, 1271–1276 (2003).

4. R. Seifert, K. Wenzel-Seifert, Constitutive activity of G-protein-coupled receptors: cause of disease and common property of wild-type receptors. Naunyn-Schmiedeberg’s Archives of Pharmacology 366, 381–416 (2002).

5. D. Calebiro, Z. Koszegi, Y. Lanoiselée, T. Miljus, S. O’Brien, G protein-coupled receptor-G protein interactions: a single-molecule perspective. Physiological Reviews 101, 857–906 (2021).

6. R. A. Cerione et al., Mammalian .beta.2-adrenergic receptor: reconstitution of functional interactions between pure receptor and pure stimulatory nucleotide binding protein of the adenylate cyclase system. Biochemistry 23, 4519–4525 (1984).

7. T. Costa, A. Herz, Antagonists with negative intrinsic activity at delta opioid receptors coupled to GTP-binding proteins. Proceedings of the National Academy of Sciences 86, 7321–7325 (1989).

8. S. T. Menon, M. Han, T. P. Sakmar, Rhodopsin: Structural Basis of Molecular Physiology. Physiol. Rev. 81, 1659–1688 (2001).

9. P. S.-H. Park, Constitutively Active Rhodopsin and Retinal Disease. Advances in pharmacology 70, 1–36 (2014).

10. H. Schihada et al., Development of a Conformational Histamine H3 Receptor Biosensor for the Synchronous Screening of Agonists and Inverse Agonists. ACS Sensors 5, 1734–1742 (2020).

11. A. L. Martin, M. A. Steurer, R. S. Aronstam, Constitutive Activity among Orphan Class-A G Protein Coupled Receptors. PLOS ONE 10, e0138463 (2015).

12. D. Araç et al., A novel evolutionarily conserved domain of cell-adhesion GPCRs mediates autoproteolysis. The EMBO Journal 31, 1364–1378 (2012).

13. T. Quon et al., Biased constitutive signaling of the G protein-coupled receptor GPR35 suppresses gut barrier permeability. Journal of Biological Chemistry 301, 108035 (2025).

14. C. Galés et al., Real-time monitoring of receptor and G-protein interactions in living cells. Nature Methods 2, 177–184 (2005).

15. C. Galés et al., Probing the activation-promoted structural rearrangements in preassembled receptor–G protein complexes. Nature Structural & Molecular Biology 13, 778–786 (2006).

16. A. S. Tyson et al., Molecular mechanisms of inverse agonism via κ-opioid receptor–G protein complexes. Nature Chemical Biology 21, 1046–1057 (2025).

17. A. Kusumi et al., Dynamic Organizing Principles of the Plasma Membrane that Regulate Signal Transduction: Commemorating the Fortieth Anniversary of Singer and Nicolson’s Fluid-Mosaic Model. Annu. Rev. Cell Dev. Biol. 28, 215–250 (2012).

18. K. Jacobson, P. Liu, B. C. Lagerholm, The Lateral Organization and Mobility of Plasma Membrane Components. Cell 177, 806–819 (2019).

19. T. Fujiwara, K. Ritchie, H. Murakoshi, K. Jacobson, A. Kusumi Phospholipids undergo hop diffusion in compartmentalized cell membrane. Journal of Cell Biology 157, 1071–1082 (2002).

20. E. Sezgin, I. Levental, S. Mayor, C. Eggeling, The mystery of membrane organization: composition, regulation and roles of lipid rafts. Nature Reviews Molecular Cell Biology 18, 361–374 (2017).

21. I. Levental, K. R. Levental, F. A. Heberle, Lipid Rafts: Controversies Resolved, Mysteries Remain. Trends Cell Biol. 30, 341–353 (2020).

22. T. Sungkaworn et al., Single-molecule imaging reveals receptor–G protein interactions at cell surface hot spots. Nature 550, 543–547 (2017).

23. K. G. N. Suzuki et al., Transient GPI-anchored protein homodimers are units for raft organization and function. Nat. Chem. Biol. 8, 774–783 (2012).

24. T. A. Spalding, E. S. Burstein, Constitutive Activity of Muscarinic Acetylcholine Receptors. Journal of Receptors and Signal Transduction 26, 61–85 (2006).

25. E. Ibrisimovic et al., Constitutive activity of the A2A adenosine receptor and compartmentalised cyclic AMP signalling fine-tune noradrenaline release. Purinergic Signalling 8, 677–692 (2012).

26. S. Keuerleber, P. Thurner, C. W. Gruber, J. Zezula, M. Freissmuth, Reengineering the Collision Coupling and Diffusion Mode of the A2A-adenosine Receptor. J. Biol. Chem. 287, 42104–42118 (2012).

27. P. Thurner et al., A two-state model for the diffusion of the A2A adenosine receptor in hippocampal neurons: agonist-induced switch to slow mobility is modified by synapse-associated protein 102 (SAP102). J Biol Chem 289, 9263–9274 (2014).

28. R. J. Ward, E. Alvarez-Curto, G. Milligan, “Using the Flp-In™ T-Rex™ System to Regulate GPCR Expression” in Receptor Signal Transduction Protocols: Third Edition, G. B. Willars, R. A. J. Challiss, Eds. (Humana Press, Totowa, NJ, 2011), 10.1007/978-1-61779-126-0_2, pp. 21–37.

29. Z. Cong et al., Molecular features of the ligand-free GLP-1R, GCGR and GIPR in complex with Gs proteins. Cell Discovery 10, 18 (2024).

30. S. A. M. Martins et al., Monitoring intracellular calcium in response to GPCR activation using thin-film silicon photodiodes with integrated fluorescence filters. Biosensors and Bioelectronics 52, 232–238 (2014).

31. Y. Li, R. V. Shivnaraine, F. Huang, J. W. Wells, C. C. Gradinaru, Ligand-Induced Coupling between Oligomers of the M_2_ Receptor and the G_i1_ Protein in Live Cells. Biophysical Journal 115, 881–895 (2018).

32. S. van de Linde et al., Photoinduced formation of reversible dye radicals and their impact on super-resolution imaging. Photochemical & Photobiological Sciences 10, 499–506 (2011).

33. P. Mystek, B. Rysiewicz, J. Gregrowicz, M. Dziedzicka-Wasylewska, A. Polit, G\gamma and G\alpha Identity Dictate a G-Protein Heterotrimer Plasma Membrane Targeting. Cells 8, 1246 (2019).

34. P. H. M. Lommerse et al., Single-Molecule Diffusion Reveals Similar Mobility for the Lck, H-Ras, and K-Ras Membrane Anchors. Biophysical Journal 91, 1090–1097 (2006).

35. F. Simon, C. H. Wiggins, L. E. Weiss, Analyzing single-molecule dynamics with both complex types of motion and complex transition kinetics: Benchmarking of ExaTrack. bioRxiv 10.64898/2026.01.22.700663 (2026).

36. P. Varshney, V. Yadav, N. Saini, Lipid rafts in immune signalling: current progress and future perspective. Immunology 149, 13–24 (2016).

37. N. Fricke et al., High-Content Imaging Platform to Discover Chemical Modulators of Plasma Membrane Rafts. ACS Central Science 8, 370–378 (2022).

38. S. K. Rodal et al., Extraction of Cholesterol with Methyl-β-Cyclodextrin Perturbs Formation of Clathrin-coated Endocytic Vesicles. Molecular Biology of the Cell 10, 961–974 (1999).

39. T. W. Sirk, E. F. Brown, A. K. Sum, M. Friedman, Molecular Dynamics Study on the Biophysical Interactions of Seven Green Tea Catechins with Lipid Bilayers of Cell Membranes. Journal of Agricultural and Food Chemistry 56, 7750–7758 (2008).

40. S. K. Huang et al., Delineating the conformational landscape of the adenosine A2A receptor during G protein coupling. Cell 10.1016/j.cell.2021.02.041 (2021).

41. M. Yanagawa et al., Single-molecule diffusion-based estimation of ligand effects on G protein–coupled receptors. Science Signaling 11, eaao1917 (2018).

42. B. K. Kobilka, X. Deupi, Conformational complexity of G-protein-coupled receptors. Trends in Pharmacological Sciences 28, 397–406 (2007).

43. S. Lu et al., Activation pathway of a G protein-coupled receptor uncovers conformational intermediates as targets for allosteric drug design. Nat Commun 12, 4721 (2021).

44. R. Lamichhane et al., Single-molecule view of basal activity and activation mechanisms of the G protein-coupled receptor β2 AR. Proceedings of the National Academy of Sciences of the United States of America 112, 14254–14259 (2015).

45. D. D. Fernandes et al., Ligand modulation of the conformational dynamics of the A2A adenosine receptor revealed by single-molecule fluorescence. Scientific Reports 11, 5910 (2021).

46. I. Maslov et al., Sub-millisecond conformational dynamics of the A2A adenosine receptor revealed by single-molecule FRET. Communications Biology 6, 362 (2023).

47. M. Klinger et al., Removal of the carboxy terminus of the A2A-adenosine receptor blunts constitutive activity: differential effect on cAMP accumulation and MAP kinase stimulation. Naunyn-Schmiedeberg’s Archives of Pharmacology 366, 287–298 (2002).

48. J. Xu et al., Conformational Complexity and Dynamics in a Muscarinic Receptor Revealed by NMR Spectroscopy. Molecular Cell 75, 53–65.e57 (2019).

49. S. Chang, E. M. Ross, Activation Biosensor for G Protein-Coupled Receptors: A FRET-Based m1 Muscarinic Activation Sensor That Regulates Gq. PLOS ONE 7, e45651 (2012).

50. N. Ziegler, J. Bätz, U. Zabel, M. J. Lohse, C. Hoffmann, FRET-based sensors for the human M1-, M3-, and M5-acetylcholine receptors. Bioorganic & Medicinal Chemistry 19, 1048–1054 (2011).

51. C. Dietrich, B. Yang, T. Fujiwara, A. Kusumi, K. Jacobson, Relationship of lipid rafts to transient confinement zones detected by single particle tracking. Biophys J 82, 274–284 (2002).

52. M. Fallahi-Sichani, J. J. Linderman, Lipid raft-mediated regulation of G-protein coupled receptor signaling by ligands which influence receptor dimerization: a computational study. PLoS One 4, e6604 (2009).

53. R. Baccouch, E. Rascol, K. Stoklosa, I. D. Alves, The role of the lipid environment in the activity of G protein coupled receptors. Biophysical Chemistry 285, 106794 (2022).

54. S. Gahbauer, R. A. Böckmann, Membrane-Mediated Oligomerization of G Protein Coupled Receptors and Its Implications for GPCR Function. Frontiers in Physiology Volume 7 - 2016 (2016).

55. A. D. Goddard, A. Watts, Regulation of G protein-coupled receptors by palmitoylation and cholesterol. BMC Biology 10, 27 (2012).

56. N. Thakur et al., Anionic phospholipids control mechanisms of GPCR-G protein recognition. Nature Communications 14, 794 (2023).

57. X. Tian, D. S. Kang, J. L. Benovic, “β-Arrestins and G Protein-Coupled Receptor Trafficking” in Arrestins - Pharmacology and Therapeutic Potential, V. V. Gurevich, Ed. (Springer Berlin Heidelberg, Berlin, Heidelberg, 2014), 10.1007/978-3-642-41199-1_9, pp. 173–186.

58. R. Janicot et al., Direct interrogation of context-dependent GPCR activity with a universal biosensor platform. Cell 187, 1527–1546.e1525 (2024).

59. X. Zhou et al., Diffusion and Oligomerization States of the Muscarinic M1 Receptor in Live Cells─The Impact of Ligands and Membrane Disruptors. The Journal of Physical Chemistry B 128, 4354–4366 (2024).

60. J. Novotny, P. Svoboda, The long (Gs(alpha)-L) and short (Gs(alpha)-S) variants of the stimulatory guanine nucleotide-binding protein. Do they behave in an identical way? Journal of Molecular Endocrinology 20, 163–173 (1998).

61. W. Stallaert et al., Purinergic Receptor Transactivation by the \beta_2-Adrenergic Receptor Increases Intracellular Ca^{2+} in Nonexcitable Cells. Mol. Pharmacol. 91, 533–544 (2017).

62. R. Schrage et al., The experimental power of FR900359 to study Gq-regulated biological processes. Nat. Commun. 6, 10156 (2015).

63. X. Zhang et al., Structural basis for the ligand recognition and signaling of free fatty acid receptors. Science Advances 10, eadj2384 (2024).

64. J.-Y. Tinevez et al., TrackMate: An open and extensible platform for single-particle tracking. Methods 115, 80–90 (2017).

65. F. Simon, J.-Y. Tinevez, S. van Teeffelen, ExTrack characterizes transition kinetics and diffusion in noisy single-particle tracks. Journal of Cell Biology 222 (2023).

66. C. B. Müller et al., Precise measurement of diffusion by multi-color dual-focus fluorescence correlation spectroscopy. Europhysics Letters 83, 46001 (2008).

67. E. Haustein, P. Schwille, Fluorescence Correlation Spectroscopy: Novel Variations of an Established Technique. Annu. Rev. Biophys. Biomol. Struct. 36, 151–169 (2007).

68. T. Bickel, A note on confined diffusion. Physica A: Statistical Mechanics and its Applications 377, 24–32 (2007).

69. A. Doicu, T. Trautmann, F. Schreier, “Tikhonov regularization for linear problems” in Numerical Regularization for Atmospheric Inverse Problems, A. Doicu, T. Trautmann, F. Schreier, Eds. (Springer Berlin Heidelberg, Berlin, Heidelberg, 2010), 10.1007/978-3-642-05439-6_3, pp. 39–106.

70. M. C. Li, Q. Q. Liu, X. Y. Lu, Y. L. Zhang, L. L. Wang, Heterologous expression of human costimulatory molecule B7-2 and construction of B7-2 immobilized polyhydroxyalkanoate nanoparticles for use as an immune activation agent. BMC Biotechnol 12, 43 (2012).

