## Supporting Information for "Local confinement within plasma membrane nanodomains promotes constitutive activity of GPCRs"

This file includes Supplementary Materials and Methods, Eqs. S1 to S23, figs. S1 to S6, and tables S1 to S4.

### SUPPLEMENTARY MATERIALS AND METHODS

#### FCS/FCCS Fitting Models

Correlation curves from FCS and FCCS measurements were analyzed using a custom MATLAB implementation of the Marquardt–Levenberg algorithm, as described previously (31). The detection-volume parameters $s$ and $w$ were obtained from independent calibration measurements using Rhodamine 6G (**fig. S1A**) and Atto655-maleimide (**fig. S1B**), with diffusion coefficients of 414 and 407 $\mu\mathrm{m}^2/s$, respectively, in aqueous phosphate-buffered saline (66). Each calibration curve was fitted using:

$$G(\tau) = G(0)\left(1+\frac{\tau}{\tau_d}\right)^{-1}\left(1+\frac{\tau}{s^2\tau_d}\right)^{-1/2}\left[1+\frac{f_t}{1-f_t}\exp\left(-\frac{\tau}{\tau_t}\right)\right]. \tag{S1}$$

The fitted aspect ratio $s$ and calculated waist $w^2 = 4D\tau_d$ were fixed for subsequent FCCS calibration and live-cell FCS measurements.

Green-red overlapping volume correction factors (OVCFs) were estimated using a self-hybridized 40-base-pair double-stranded DNA labelled with Cy3 and Cy5 at opposite ends (**fig. S1C**). The primary strand sequence was 5’-TA AGC CTC GTC CTG CGT CGG AGC CCG TCT GCC AGC GGA AT-3’. The green and red autocorrelation curves were fitted using **Eq. S1**, and the cross-correlation curve was fitted using:

$$G_x(\tau) = G_x(0)\left(1+\frac{\tau}{\tau_d}\right)^{-1}\left(1+\frac{\tau}{s^2\tau_d}\right)^{-1/2}. \tag{S2}$$

For an ideal FCCS positive control, the green and red autocorrelation amplitudes and the cross-correlation amplitude are identical. Deviations were assigned to non-ideal overlap between the two detection volumes, and OVCFs were calculated as:

$$\begin{aligned}\mathrm{OVCF}_g &= \frac{G_r(0)}{G_x(0)},\\ \mathrm{OVCF}_r &= \frac{G_g(0)}{G_x(0)}.\end{aligned} \tag{S3}$$

For single-color FCS of GPCRs, bottom-membrane intensity fluctuations were fitted with a two-component 2D diffusion model and a photophysical dark state (67):

$$\begin{aligned}G(\tau) = &\frac{1}{\langle N\rangle}\left[f_1\left(1+\frac{\tau}{\tau_{D1}}\right)^{-1} + (1-f_1)\left(1+\frac{\tau}{\tau_{D2}}\right)^{-1}\right]\\ &\times\left[1+\frac{f_{ds}}{1-f_{ds}}\exp\left(-\frac{\tau}{\tau_{ds}}\right)\right].\end{aligned} \tag{S4}$$

Here, $\tau_{D1}$ and $\tau_{D2}$ are the residence times of the two diffusing species, $f_1$ is the fraction of the first diffusion component, $\langle N\rangle$ is the average number of fluorescent molecules in the detection volume, and $\tau_{ds}$ and $f_{ds}$ are the lifetime and population fraction of the fluorophore dark state. Diffusion coefficients were calculated from $D = w_0^2/(4\tau_D)$, where $w_0$ is the lateral waist of the confocal detection volume. Correlation curves were fitted from 100 $\mu$s to 1 s to focus on membrane diffusion while excluding faster photophysical dynamics.

For single-color FCS of G proteins, a one-component 2D diffusion model with the same dark-state term was sufficient:

$$G(\tau) = \frac{1}{\langle N \rangle}\left(1 + \frac{\tau}{\tau_D}\right)^{-1}\left[1 + \frac{f_{ds}}{1 - f_{ds}}\exp\left(-\frac{\tau}{\tau_{ds}}\right)\right]. \qquad \text{(S5)}$$

For FCCS, the green-channel GPCR and red-channel G protein autocorrelation curves were fitted globally using **Eq. S4** with a shared slow diffusion coefficient $D_2$. Because the dark-state dynamics of JF549i-HTL and JF669-STL attached to different proteins were assumed to be uncorrelated, the green-red cross-correlation curve was fitted using:

$$G_x(\tau) = \frac{\langle N_x \rangle}{\langle N_g \rangle \langle N_r \rangle}\left(1 + \frac{\tau}{\tau_{Dx}}\right)^{-1}. \qquad \text{(S6)}$$

Here, $N_g$ and $N_r$ are the average numbers of green and red fluorescent molecules in the overlapping detection volume, $N_x$ is the average number of codiffusing species, and $\tau_{Dx}$ is their average residence time. The codiffusion coefficient $D_x = w_0^2/(4\tau_{Dx})$ was globally shared as $D_2$ in the autocorrelation curves. The degree of GPCR–G protein association was estimated as the fraction of each fluorescent species that codiffuses with the other:

$$\begin{aligned} fcd_g &= \frac{\langle N_x \rangle}{\langle N_g \rangle}\,\mathrm{OVCF}_g, \\ fcd_r &= \frac{\langle N_x \rangle}{\langle N_r \rangle}\,\mathrm{OVCF}_r. \end{aligned} \qquad \text{(S7)}$$

Associations between $fcd_r$ and receptor surface density were assessed using two-sided Spearman rank correlation tests.

### SINGLE-PARTICLE TRACKING AND EXATRACK ANALYSIS

Particle positions and intensities in each TIRF frame were obtained with the difference-of-Gaussian detector in TrackMate, and two-dimensional trajectories were assembled using the Linear Assignment Problem tracker (59). For GPCR tracking, a 0.33 $\mu$m linking radius and a maximum gap of 60 ms were used. The maximum linking distance was selected by modelling Brownian step lengths $\Delta r$ with a Rayleigh distribution:

$$P(\Delta r) = \frac{\Delta r}{\sigma^2}\exp\left(-\frac{\Delta r^2}{2\sigma^2}\right), \qquad \sigma^2 = 2D\Delta t, \qquad \Delta r_{98} \approx 3\sigma. \qquad \text{(S8)}$$

An initial tracking pass used $\Delta r_{98}$ as the maximum linking distance. The resulting step-size distribution was compared with the nearest-neighbour half-distance distribution of all detected particles. The final linking distance was not smaller than $\Delta r_{98}$, ensuring coverage of expected displacements, and not larger than the fifth percentile of the nearest-neighbour half-distance distribution, limiting mis-linking in crowded regions.

Gap-closing parameters were calibrated from fluorophore blinking in fixed-cell movies. Reference particle positions were identified from the summed intensity of the first 20 frames, excluding particles within one particle diameter of one another. Localizations were assigned to the nearest reference position within one pixel, and blinking events were defined as intervals lacking an assigned localization. The maximum frame gap was set from the median blink duration. For a particle with maximum diffusion coefficient $D_{\max}$ and frame gap $\Delta t_{\text{gap}}$, the theoretical gap-closing distance was:

$$\Delta r_{\text{gap}} = 3\sigma_{\text{gap}} = 3\sqrt{2D_{\max}\Delta t_{\text{gap}}}. \tag{S9}$$

This bound was empirically tested over a range of gap-closing distances, and the final value was selected at the inflection point in track count or mean track length.

Individual time-averaged mean-squared displacement curves were calculated as:

$$\text{MSD}(\tau) = \frac{1}{T-\tau}\sum_{t=0}^{T-\tau}\{[x(t+\tau)-x(t)]^2 + [y(t+\tau)-y(t)]^2\}. \tag{S10}$$

Ensemble time-averaged MSD curves were obtained as variance-weighted averages of the individual curves. Trajectories classified by ExaTrack as $\geq 80\%$ Brownian were fitted with:

$$\text{MSD}(\tau) = 4D\tau + \sigma^2. \tag{S11}$$

where $\sigma^2$ represents localization precision. Trajectories classified as $\geq 80\%$ confined diffusion were fitted with:

$$\text{MSD}(\tau) = a^2\left[1 - 8\sum_{m=1}^{\infty}\frac{\exp(-\alpha_{1m}^2\tau D/a^2)}{\alpha_{1m}^2(\alpha_{1m}^2-1)}\right] + \sigma^2. \tag{S12}$$

where $\alpha_{1m}$ is the $m^{th}$ positive root of the Bessel function of the first kind (68), and $a$ is the confinement radius.

For a confined segment in ExaTrack, the particle was modelled within a quadratic potential well whose center itself diffuses. For particle positions $r_i$ and potential-well centers $h_i$, the joint probability density was:

$$\begin{gathered} f_u(r_0)f_v(h_0 - r_0)\prod_{i=0}^{n} f_\sigma(r_i - c_i) \\ \times f_d[r_{i+1} - (1-l)r_i - lh_i]f_q(h_{i+1} - h_i). \end{gathered} \tag{S13}$$

Here, $f_y(x)$ denotes a zero-mean Gaussian density with standard deviation $y$, $c_i$ is the observed localization, and the confinement factor $l$ ranges from zero for free diffusion to one for complete attachment to the potential-well center (35). Integration over particle positions, well centers, and possible state sequences gives the track likelihood used for maximum-likelihood estimation. The confinement radius containing 95% of localizations was calculated as:

$$R_{\text{conf}} = 2\sqrt{\frac{D\Delta t}{l}}. \tag{S14}$$

Candidate models with two to six physical diffusion states were fitted with ExaTrack and compared using a regularized Bayesian information criterion:

$$\begin{aligned} \mathrm{BIC} &= -2\ell + k\ln n, \\ \mathrm{BIC}_{\mathrm{reg}} &= \mathrm{BIC} + \lambda k \ell, \qquad \lambda = 0.0002. \end{aligned} \tag{S15}$$

Here, $\ell$ is the fitted log-likelihood, $k$ is the number of free model parameters, and $n$ is the number of observations. Lower $\mathrm{BIC}_{\mathrm{reg}}$ values indicate a preferred statistical description; the final state number was also assessed from the fitted diffusion coefficients, state lifetimes, and physical interpretation of the states.

For localization $t$ in trajectory $k$, the continuous diffusion-state index used for spatial maps was calculated from ExaTrack posterior probabilities using **Eq. 1** in the main text, where $p_i(k,t)$ is the posterior probability of state $i$. State-index values were then averaged over neighbouring localizations within a one-pixel radius (0.167 $\mu$m) to obtain $S'(\mathbf{r})$, as defined in **Eq. 2**.

Diffusion maps were plotted with each localization coloured by $S'(\mathbf{r})$, normalized from 1 to 3 to represent immobile, confined, and free/Brownian diffusion. Cell boundaries from the underlying TIRF image were overlaid as black contours.

### IMAGE CROSS-CORRELATION SPECTROSCOPY

Image cross-correlation spectroscopy (ICCS) was used to quantify spatial overlap between receptor and G protein confinement domains. The input for each spectral channel was the time-averaged diffusion-state map $S'(\mathbf{r})$ defined in **Eq. 2**, restricted to the cell contour. For co-confinement analysis, positions with $S'(\mathbf{r}) < 2.25$ were retained to emphasize immobile and confined states on the 1–3 diffusion-state scale. The resulting map signal in channel $a$ was denoted $M_a(\mathbf{r})$, and its mean-subtracted fluctuation was:

$$\delta M_a(\mathbf{r}) := M_a(\mathbf{r}) - \langle M_a(\mathbf{r}) \rangle. \tag{S16}$$

Because the diffusion maps were time averaged, ICCS was evaluated only as a function of spatial displacement. The normalized spatial covariance between channels $a$ and $b$ was calculated as:

$$g_{ab}(\Delta\mathbf{r}) = \frac{\langle \delta M_a(\mathbf{r}+\Delta\mathbf{r}) \delta M_b(\mathbf{r}) \rangle_{\mathbf{r}}}{\sqrt{\langle \delta M_a^2(\mathbf{r}) \rangle_{\mathbf{r}} \langle \delta M_b^2(\mathbf{r}) \rangle_{\mathbf{r}}}}. \tag{S17}$$

Radial averaging converted the two-dimensional covariance into a one-dimensional spatial correlation profile,

$$G_{ab}(r) = \langle g_{ab}(\Delta\mathbf{r}) \rangle_{\|\Delta\mathbf{r}\|=r}. \tag{S18}$$

where autocorrelation curves $G_{aa}(r)$ and $G_{bb}(r)$ report the characteristic spatial scale of confinement domains in each channel, and the cross-correlation curve $G_{ab}(r)$ reports their overlap. Measured radial autocorrelation curves were broadened by finite localization precision, the optical point spread function, and spatial averaging during map generation. For each channel, the measured autocorrelation $y(r)$ was therefore modeled as the convolution of an interaction-only signal $g(r)$ with a simulated point-spread-function baseline $k(r)$, plus a residual term $\varepsilon(r)$:

$$y(r) \approx (k * g)(r) + \varepsilon(r). \tag{S19}$$

$g(r)$ was estimated by Wiener–Tikhonov deconvolution filtering with zero-padding to limit circular wrap-around (69):

$$\begin{aligned} \hat{g}_\lambda(\omega) &= \frac{K^*(\omega)}{|K(\omega)|^2 + \lambda} Y(\omega), \\ g_\lambda &= \mathcal{F}^{-1}\{\hat{g}_\lambda\}. \end{aligned} \tag{S20}$$

where $Y = \mathcal{F}\{y\}$ and $K = \mathcal{F}\{k\}$, with $\mathcal{F}$ denoting the two-dimensional, real-to-complex Fourier transform and $^*$ denoting complex conjugation. The scalar $\lambda$ regularizes the ill-posed inverse by trading bias for variance.

After deconvolution, the short-range amplitudes of the corrected receptor autocorrelation function, $g_R(0)$, and the receptor–G protein cross-correlation function, $g_x(0)$, were estimated as the arithmetic mean of correlation values at radial lags $r < 10\ px$. These amplitudes were then used to calculate the fraction of co-confinement (*coco*) using **Eq. 3** in the main text. Associations between co-confinement and apparent receptor or G protein density were assessed using two-sided Spearman rank correlation tests.

**Mean Encounter Rates**

To estimate how membrane confinement changes receptor–G protein encounter kinetics, we used the two-dimensional Smoluchowski approximation for diffusion-limited reactions. The relative diffusion coefficient of the receptor and G protein was calculated as:

$$D_{\text{rel}} = D_R + D_G. \tag{S21}$$

where $D_R$ and $D_G$ are the receptor and G protein diffusion coefficients, respectively. The corresponding two-dimensional bimolecular encounter-rate constant was:

$$k_{2D} = \frac{2\pi D_{\text{rel}}}{\ln(L/a) - \gamma}. \tag{S22}$$

Here, $a \approx 8$ nm is the capture radius, $L$ is the characteristic search length, and $\gamma \approx 0.577$ is the Euler–Mascheroni constant. The pseudo-first-order encounter rate for one receptor was then calculated from the local G protein surface density, $\rho_G$, as:

$$k_{\text{enc}} = k_{2D}\rho_G. \tag{S23}$$

For the cell-scale estimate, $L \approx 6.5\ \mu\text{m}$ and $\rho_G \approx 0.1$ molecules/$\mu\text{m}^2$, giving $k_{\text{enc}} \approx 0.075\ \text{s}^{-1}$, or a mean encounter time of approximately 13.3 s. For a confinement domain, $L \approx 160$ nm and the local density was estimated as one G protein within a circular domain of this diameter, giving $k_{\text{enc}} \approx 5.22\ \text{s}^{-1}$, or a mean encounter time of approximately 0.19 s. These estimates quantify the kinetic effect of local concentration within a confinement domain and do not assume stable receptor–G protein pre-coupling.

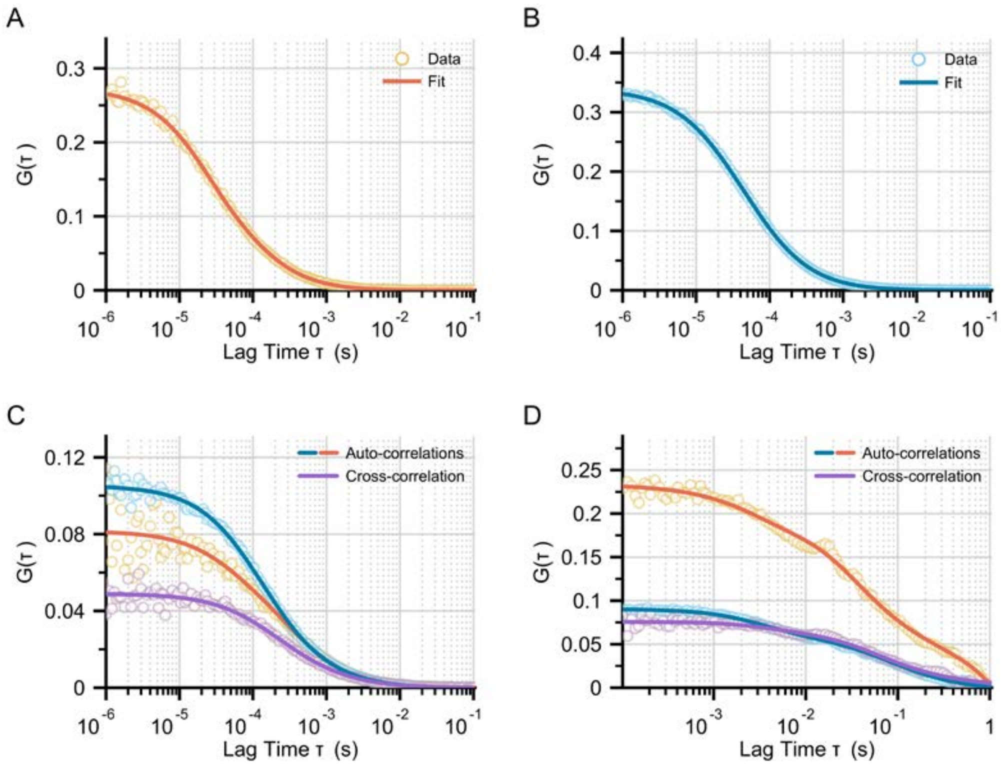


**Figure S1. Calibration FCS and FCCS measurements.** Representative calibration curves are shown in **(A–C)**; uncertainties are from the calibration fits. Dyes in **(A,B)** were measured in aqueous phosphate-buffered saline. FCS Detection volume parameters were estimated using Rhodamine 6G for the green channel **(A;** $w = 321.2 \pm 2.3\ nm$**,** $s = 6.68 \pm 0.17$**)** and Atto655-maleimide for the red channel **(B;** $w = 280.0 \pm 1.1\ nm$**,** $s = 7.96 \pm 0.28$**)**. **(C)** FCCS calibration was performed using Cy3 and Cy5 labelled 40-base-pair double-strand DNA, which yielded overlapping volume correction factors of $OVCF_g = 2.14 \pm 0.02$ and $OVCF_r = 1.40 \pm 0.01$ assuming $fcd = 1$ (**Eq. S3**). **(D)** Live-cell FCCS was tested in Flp-In T-REx 293 cells stably expressing monomeric SNAP-Halo-CD86 (70); the red-channel autocorrelation reports SNAPf-Tag labelled with JF669-STL, and the green-channel autocorrelation reports HaloTag labelled with JF549i-HTL. Estimated co-diffusion fractions using **Eq. S7** were $fcd_r = 0.54 \pm 0.01$ and $fcd_g = 0.97 \pm 0.02$. Values in **(D)** are mean $\pm$ SEM from *n* = 3 cells in one biological replicate.

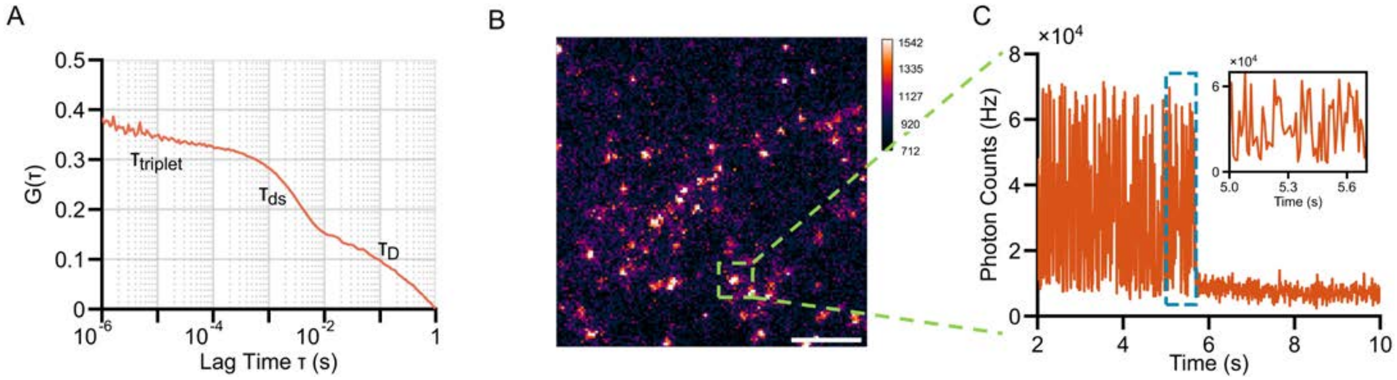


**Figure S2. The nature of the millisecond kinetics in FCS measurements. (A)** Complete raw autocorrelation (AC) curve of Halo-$A_{2A}$R labelled with JF549i-HTL in a live Flp-In T-REx 293 cell stably expressing Halo-$A_{2A}$R. Decays at different timescales are observed, with the fastest component on the microsecond scale corresponding to the triplet-state lifetime of the fluorophore. The AC curve next shows a decay component on a low-millisecond scale, which corresponds to a very fast diffusion coefficient of $\sim 5\ \mu m^2/s$. **(B)** Selected frame from a TIRF image series of fixed Flp-In T-REx 293 cells stably expressing Halo-$A_{2A}$R labelled with JF549i-HTL, acquired at 100 fps using an excitation intensity comparable to FCS experiments, $\sim 1\ kW/cm^2$; scale bar: 5 $\mu m$. **(C)** Intensity trace of a typical fluorescent spot from the TIRF image series, showing millisecond-scale photoblinking before the final photobleaching step.

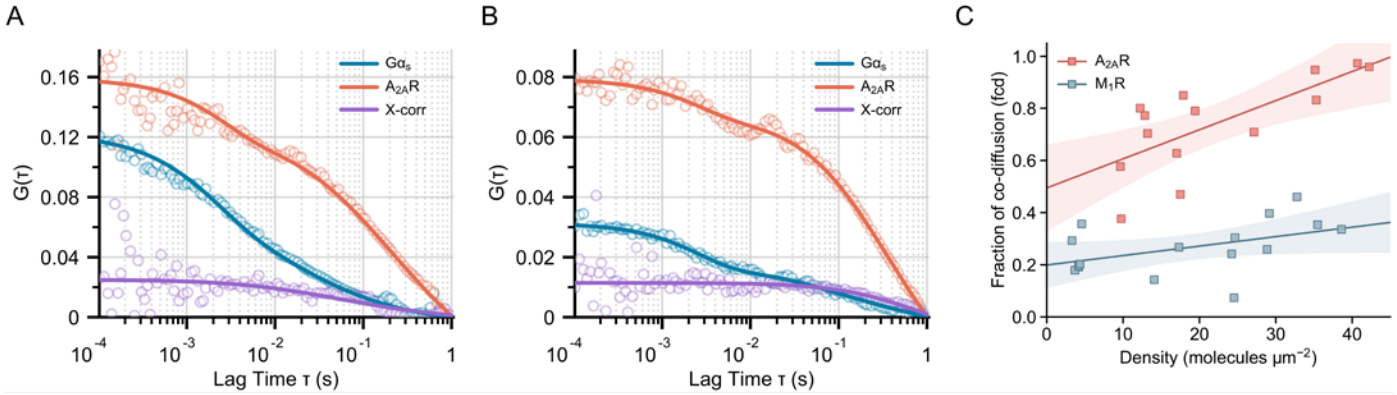


**Figure S3. Density dependence of GPCR–G protein co-diffusion.** Representative apo state (no ligand) FCCS decays for $A_{2A}R$ and $G\alpha_s$ measured at lower **(A)** and higher **(B)** brightness regions in live cells: AC of receptors (orange); AC of G proteins (cyan); X-corr of receptors and G proteins (violet). Estimated local densities of receptors were $17.5 \pm 0.7$ molecules/$\mu m^2$ **(A)** and $35.3 \pm 2.8$ molecules/$\mu m^2$ **(B)**, while the corresponding G protein densities were $67.5 \pm 3.7$ molecules/$\mu m^2$ **(A)** and $108.9 \pm 9.7$ molecules/$\mu m^2$ **(B)**. Density values in **(A,B)** are mean ± SEM from measurements in one cell. Approximately 47% of $A_{2A}R$ co-diffused with $G\alpha_s$ in **(A)**, compared with $\sim$ 83% in **(B)**. **(C)** Fraction of $M_1R$ or $A_{2A}R$ co-diffusing with $G\alpha_{11}$ or $G\alpha_s$, respectively, as a function of receptor density in the apo state. Lines show linear fits and shaded regions indicate 95% confidence bands. Two-sided Spearman rank correlation tests gave $\rho = 0.55$, $P = 0.027$ for $M_1R$ and $\rho = 0.75$, $P = 0.002$ for $A_{2A}R$. FCCS used stable $G\alpha_{11}$-SNAP or $G\alpha_s$-SNAP Flp-In T-REx 293 cells transiently expressing the cognate Halo-GPCR. Panels **(A–C)** are from the apo datasets in **Table 2**: *n* = 16 cells for $M_1R$–$G\alpha_{11}$ and *n* = 14 cells for $A_{2A}R$–$G\alpha_s$, collected across three independent biological replicates per condition. Each point in **(C)** represents one cell.

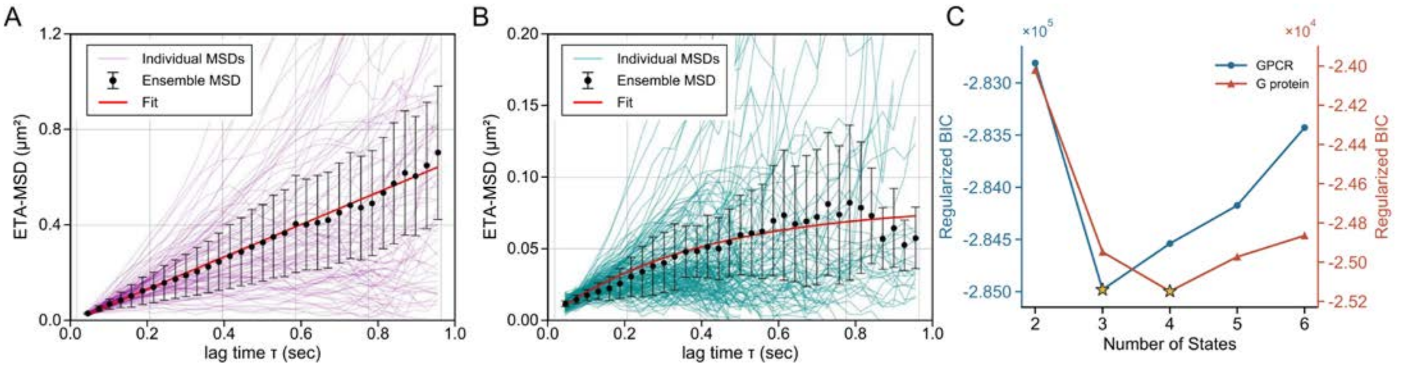


**Figure S4. Ensemble time-averaged mean-squared displacement (ETA-MSD) and ExaTrack model justification. (A)** ETA-MSD for $A_{2A}R$ undergoing Brownian motion, weighted fit with **Eq. S11** resulting in a diffusion coefficient of $D = 0.161 \pm 0.005\ \mu m^2/s$ and localization precision of $\sigma = 35 \pm 3\ nm$. **(B)** ETA-MSD for $A_{2A}R$ undergoing confined diffusion, weighted fit with **Eq. S12** resulting in $D = 0.051 \pm 0.004\ \mu m^2/s$, $\sigma = 32 \pm 4\ nm$, and a confinement radius of $a = 155 \pm 10\ nm$. **(C)** Regularized BIC ($\lambda = 0.0002$; **Eq. S15**) for two- to six-state ExaTrack models fitted to representative GPCR ($A_{2A}R$; blue circles, left axis) and G-protein ($G\alpha_s$; red triangles, right axis) datasets. Gold stars mark the minima at three states for GPCR and four states for G protein. Panels **(A–C)** show representative SPT analyses from *n* = 1 Flp-In T-REx 293 cell expressing $A_{2A}R$ and $G\alpha_s$. In **(A,B)**, ensemble curves are variance-weighted averages of individual time-averaged MSD curves; error bars show the weighted SD at each lag time. The ± values for fitted parameters in **(A,B)** denote fitting errors.

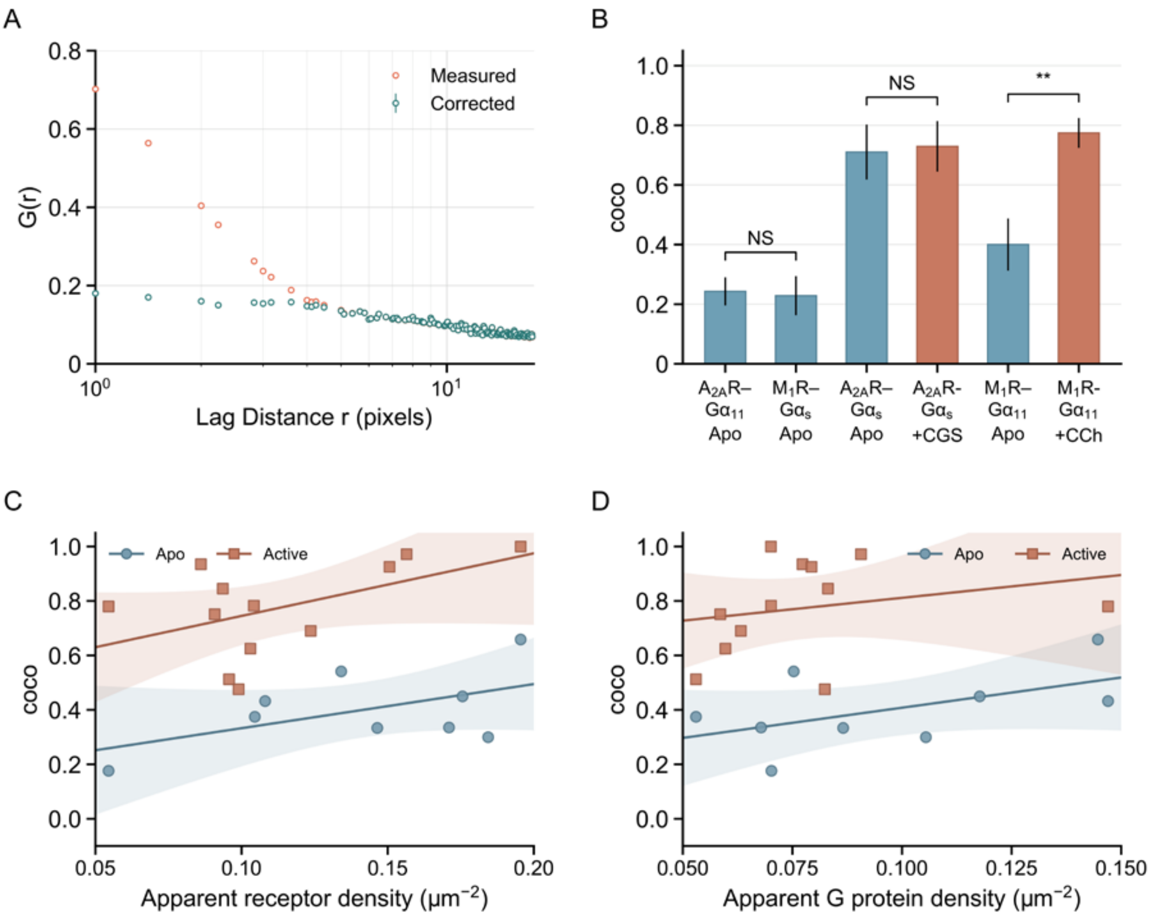


**Figure S5. ICCS correction, co-confinement controls, and density dependence.** **(A)** Representative raw (orange) and corrected (turquoise) AC decay after deconvolving the contribution of the point spread function, from the same apo $A_{2A}R$–$G\alpha_s$ example cell shown in Fig. 6I (selected from *n* = 18 cells across three independent biological replicates). **(B)** GPCR–G protein co-confinement fractions ($coco$) under control and cognate-pair conditions. Data are mean ± SEM; cells were collected across three independent biological replicates per condition. $A_{2A}R$–$G\alpha_{11}$ and $M_1R$–$G\alpha_s$ were measured as negative cross-talk controls (*n* = 9 cells per pair). $A_{2A}R$–$G\alpha_s$ measurements in apo and agonist-stimulated states were performed in $\Delta G_s$ HEK293 cells (*n* = 10 apo and 7 agonist-stimulated cells), whereas $M_1R$–$G\alpha_{11}$ measurements were performed in $\Delta G_{q/11}$ HEK293 cells (*n* = 9 apo and 12 agonist-stimulated cells). Both tagged proteins were transiently co-expressed in the G-protein-depleted cells. Detailed SPT results in G protein depletion cells are listed in **table S4**, and $coco$ values are listed in **table S2**. **, $P < 0.01$; NS, not significant, tested by two-sided Welch's $t$-tests with Holm correction. **(C, D)** $M_1R$–$G\alpha_{11}$ $coco$ values in $\Delta G_{q/11}$ cells as a function of apparent receptor density **(C)** and apparent G protein density **(D)** under apo and agonist-stimulated conditions (*n* = 9 and 12 cells from the corresponding conditions in B, respectively; each point represents one cell). Lines show linear fits and shaded regions indicate 95% confidence bands. Two-sided Spearman rank correlation tests for receptor density gave $\rho = 0.35$, $P = 0.356$ for apo and $\rho = 0.35$, $P = 0.265$ for active conditions; for G protein density, $\rho = 0.42$, $P = 0.265$ for apo and $\rho = 0.39$, $P = 0.208$ for active conditions.

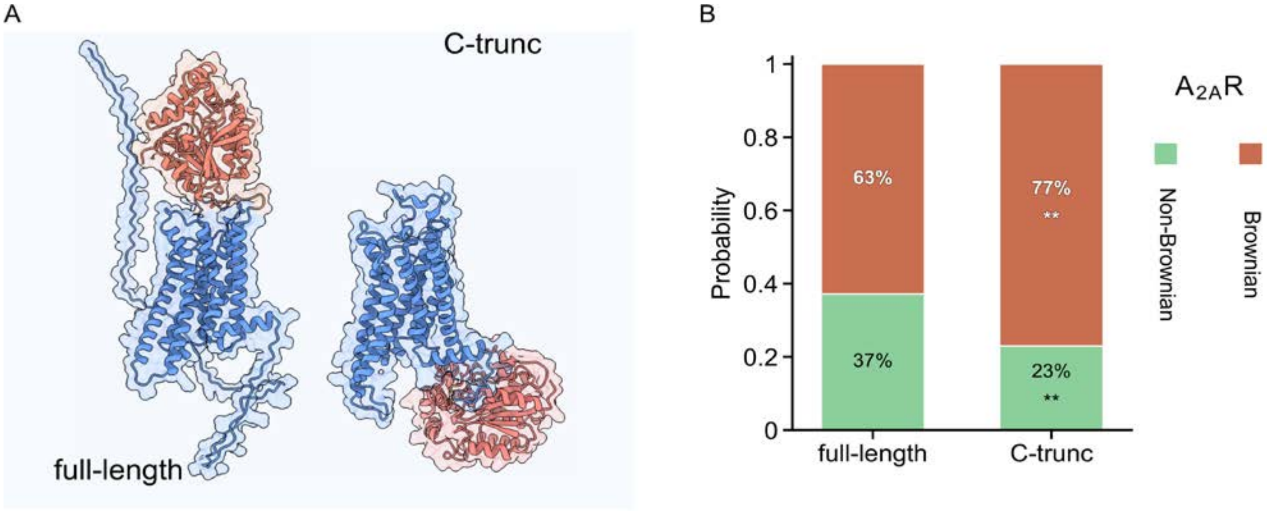


**Figure S6. C-terminal truncation reduces $A_{2A}$R confinement.** **(A)** AlphaFold-predicted structures of full-length Halo-$A_{2A}$R and the C-terminally truncated $A_{2A}$R(1–316)–Halo construct (C-trunc), with HaloTag (*red*) and $A_{2A}$R (*blue*) indicated. **(B)** Single-particle tracking (SPT) analysis of full-length and C-trunc $A_{2A}$R in Flp-In T-REx 293 cells under basal conditions. Stacked composition bars summarize the mean diffusion populations as Brownian and non-Brownian states, **, $P < 0.01$, tested by two-sided Welch's $t$-test with Holm correction. Full SPT fitting results are listed in **table S1**. Data in (B) were obtained from *n* = 23 cells expressing full-length $A_{2A}$R and *n* = 7 cells expressing C-terminally truncated $A_{2A}$R, collected across three independent biological replicates.

**Table S1.** Detailed ExaTrack analysis of SPT data for GPCRs and G proteins[a].

| Protein | Condition | $D_1$ ($\mu m^2/s$) | $f_1$ (%) | $l_1$ | $D_2$ ($\mu m^2/s$) | $f_2$ (%) | $l_2$ | $D_3$ ($\mu m^2/s$) | $l_3$ | $N$ ($\times 10^5$) | $n$ (cells) |
|---|---|---|---|---|---|---|---|---|---|---|---|
| $M_1R$ | Apo | 0.22 ± 0.02 | 75 ± 4 | 0.03 | 0.092 ± 0.006 | 20 ± 5 | 0.43 | 0.003 | 0.42 | 2.7 | 25 |
| | + MBCD | 0.17 ± 0.01 | 79 ± 2 | 0.03 | 0.054 ± 0.008 | 18 ± 2 | 0.42 | 0.003 | 0.28 | 1.0 | 12 |
| | + EGCG | 0.18 ± 0.01 | 49 ± 6 | 0.02 | 0.028 ± 0.002 | 35 ± 7 | 0.24 | 0.002 | 0.33 | 1.4 | 14 |
| | + Carbachol | 0.22 ± 0.02 | 40 ± 2 | 0.05 | 0.064 ± 0.006 | 31 ± 2 | 0.15 | 0.007 | 0.08 | 1.6 | 15 |
| $G\alpha_{11}$ | Apo | 0.55 ± 0.03 | 64 ± 8 | 0.04 | 0.059 ± 0.007 | 24 ± 7 | 0.35 | 0.001 | 0.45 | 0.8 | 15 |
| | + Carbachol | 0.63 ± 0.05 | 48 ± 6 | 0.01 | 0.120 ± 0.015 | 33 ± 5 | 0.22 | 0.008 | 0.50 | 0.6 | 13 |
| $A_{2A}R$ | Apo | 0.21 ± 0.01 | 63 ± 4 | 0.01 | 0.125 ± 0.009 | 31 ± 4 | 0.53 | 0.002 | 0.38 | 2.2 | 23 |
| | + MBCD | 0.17 ± 0.01 | 71 ± 2 | 0.06 | 0.053 ± 0.003 | 23 ± 2 | 0.24 | 0.014 | 0.28 | 0.8 | 11 |
| | + EGCG | 0.15 ± 0.01 | 29 ± 7 | 0.05 | 0.020 ± 0.002 | 40 ± 8 | 0.27 | 0.003 | 0.15 | 1.1 | 15 |
| | + CGS-21680 | 0.21 ± 0.01 | 60 ± 3 | 0.02 | 0.077 ± 0.008 | 31 ± 3 | 0.40 | 0.004 | 0.30 | 1.4 | 12 |
| | C-trunc[b] | 0.22 ± 0.01 | 77 ± 1 | 0.01 | 0.081 ± 0.005 | 20 ± 1 | 0.18 | 0.012 | 0.34 | 0.7 | 7 |
| $G\alpha_s$ | Apo | 0.45 ± 0.02 | 50 ± 4 | 0.01 | 0.088 ± 0.010 | 36 ± 6 | 0.40 | 0.001 | 0.24 | 0.9 | 18 |
| | + CGS-21680 | 0.47 ± 0.04 | 54 ± 2 | 0.02 | 0.064 ± 0.006 | 36 ± 5 | 0.21 | 0.001 | 0.22 | 0.7 | 13 |

[a] For each condition, values represent global fit averages. Parametric errors are shown as $\pm$ SEM; errors for all confinement factors ($l$) are $\pm 0.01$; errors for immobile-population diffusion coefficients ($D_3$) are $\pm 0.001$. The immobile fraction can be calculated as $f_3 = 1 - f_1 - f_2$. $l_1$, $l_2$, and $l_3$ denote confinement factors for the Brownian, confined, and immobile populations, respectively.
[b] C-trunc denotes the $A_{2A}R$(1–316)–Halo construct.

**Table S2.** Image Cross-Correlation Analysis.[a]

| Protein pair | Condition | co-confinement ($coco$) | $\boldsymbol{n}$ (cells) |
|---|---|---|---|
| $M_1R$–$G\alpha_{11}$ | Apo | $0.31 \pm 0.03$ | 15 |
| | + Carbachol | $0.52 \pm 0.05$ | 13 |
| $A_{2A}R$–$G\alpha_s$ | Apo | $0.55 \pm 0.07$ | 18 |
| | + CGS-21680 | $0.51 \pm 0.08$ | 13 |
| $M_1R$–CD59 | Apo | $0.42 \pm 0.09$ | 9 |
| | + Carbachol | $0.60 \pm 0.11$ | 11 |
| $A_{2A}R$–CD59 | Apo | $0.78 \pm 0.10$ | 8 |
| | + CGS-21680 | $0.73 \pm 0.07$ | 8 |
| $A_{2A}R$–$G\alpha_s$ | Apo, $\Delta G_s$ cells | $0.71 \pm 0.09$ | 10 |
| | + CGS-21680, $\Delta G_s$ cells | $0.73 \pm 0.08$ | 7 |
| $M_1R$–$G\alpha_{11}$ | Apo, $\Delta G_{q/11}$ cells | $0.40 \pm 0.09$ | 9 |
| | + Carbachol, $\Delta G_{q/11}$ cells | $0.77 \pm 0.05$ | 12 |
| $A_{2A}R$–$G\alpha_{11}$ | Apo | $0.24 \pm 0.05$ | 9 |
| $M_1R$–$G\alpha_s$ | Apo | $0.23 \pm 0.07$ | 9 |

[a] Co-confinement ($coco$) was calculated from the ICCS amplitudes using **Eq. 3**; errors are SEM. $\Delta G_s$ and $\Delta G_{q/11}$ denote HEK293 cell lines lacking the indicated endogenous G-protein family, in which the tagged receptor and tagged G protein were transiently co-expressed.

**Table S3.** Detailed ExaTrack analysis of SPT data for CD59[a].

| **Protein** | **Condition** | $\boldsymbol{D_1}$ ($\mu m^2/s$) | $\boldsymbol{f_1}$ (%) | $\boldsymbol{l_1}$ | $\boldsymbol{D_2}$ ($\mu m^2/s$) | $\boldsymbol{f_2}$ (%) | $\boldsymbol{l_2}$ | $\boldsymbol{D_3}$ ($\mu m^2/s$) | $\boldsymbol{l_3}$ | $\boldsymbol{N}$ ($\times 10^5$) | $\boldsymbol{n}$ (cells) |
|---|---|---|---|---|---|---|---|---|---|---|---|
| CD59 | Apo | 0.22 ± 0.01 | 62 ± 7 | 0.01 | 0.066 ± 0.037 | 23 ± 6 | 0.35 | 0.009 | 0.27 | 3.0 | 14 |
| | + MBCD | 0.23 ± 0.01 | 70 ± 3 | 0.00 | 0.075 ± 0.014 | 23 ± 3 | 0.17 | 0.003 | 0.22 | 3.7 | 17 |
| | + EGCG | 0.20 ± 0.02 | 41 ± 9 | 0.05 | 0.065 ± 0.019 | 40 ± 6 | 0.42 | 0.012 | 0.29 | 2.0 | 14 |
| | + Carbachol | 0.23 ± 0.01 | 60 ± 11 | 0.01 | 0.086 ± 0.024 | 26 ± 7 | 0.33 | 0.014 | 0.22 | 0.8 | 11 |
| | + CGS-21680 | 0.24 ± 0.01 | 64 ± 7 | 0.01 | 0.083 ± 0.024 | 25 ± 6 | 0.44 | 0.006 | 0.16 | 0.4 | 8 |

[a] For each condition, values represent weighted means. Where shown, errors are $\pm$ SEM. $D_1$, $D_2$, and $D_3$ denote the Brownian, confined, and immobile states, respectively; $f$ denotes the corresponding state fraction, $l$ the confinement factor, $N$ the number of tracks, and $n$ the number of cells.

**Table S4.** Detailed ExaTrack analysis of SPT data in delta-G cell lines for GPCRs and G proteins[a].

| **Protein** | **Ligand** | $D_1$ ($\mu m^2/s$) | $f_1$ (%) | $l_1$ | $D_2$ ($\mu m^2/s$) | $f_2$ (%) | $l_2$ | $D_3$ ($\mu m^2/s$) | $l_3$ | $N$ ($\times 10^5$) | $n$ (cells) |
|---|---|---|---|---|---|---|---|---|---|---|---|
| $M_1R$ | Apo | 0.22 ± 0.01 | 68 ± 7 | 0.01 | 0.087 ± 0.023 | 24 ± 6 | 0.29 | 0.013 | 0.29 | 0.3 | 9 |
| | + Carbachol | 0.23 ± 0.01 | 52 ± 6 | 0.01 | 0.095 ± 0.025 | 29 ± 3 | 0.35 | 0.018 | 0.17 | 0.7 | 12 |
| $G\alpha_{11}$ | Apo | 0.52 ± 0.06 | 51 ± 5 | 0.04 | 0.094 ± 0.078 | 32 ± 6 | 0.43 | 0.009 | 0.30 | 0.1 | 9 |
| | + Carbachol | 0.59 ± 0.06 | 60 ± 11 | 0.05 | 0.108 ± 0.050 | 25 ± 6 | 0.43 | 0.004 | 0.16 | 0.2 | 12 |
| $A_{2A}R$ | Apo | 0.22 ± 0.01 | 40 ± 3 | 0.02 | 0.099 ± 0.038 | 29 ± 3 | 0.65 | 0.041 | 0.27 | 0.6 | 10 |
| | + CGS-21680 | 0.22 ± 0.01 | 41 ± 8 | 0.07 | 0.084 ± 0.028 | 35 ± 5 | 0.40 | 0.032 | 0.22 | 0.8 | 7 |
| $G\alpha_s$ | Apo | 0.40 ± 0.05 | 31 ± 8 | 0.04 | 0.058 ± 0.020 | 44 ± 4 | 0.24 | 0.002 | 0.23 | 0.1 | 10 |
| | + CGS-21680 | 0.38 ± 0.05 | 31 ± 5 | 0.02 | 0.050 ± 0.015 | 44 ± 1 | 0.29 | 0.003 | 0.18 | 0.1 | 7 |

[a] $A_{2A}R$ and $G\alpha_s$ were measured in $\Delta G_s$ HEK293 cells, whereas $M_1R$ and $G\alpha_{11}$ were measured in $\Delta G_{q/11}$ HEK293 cells. For each condition, values represent weighted means. Where shown, errors are ± SEM. $D_1$, $D_2$, and $D_3$ denote the Brownian, confined, and immobile states, respectively; $f$ denotes the corresponding state fraction, $l$ the confinement factor, $N$ the number of tracks, and $n$
